# The secreted Nimrod NimB1 negatively regulates early steps of apoptotic cell phagocytosis in *Drosophila*

**DOI:** 10.1101/2025.01.07.631694

**Authors:** Asya Dolgikh, Samuel Rommelaere, Aseel Ghanem, Bianca Petrignani, Mickael Poidevin, Estee Kurant, Bruno Lemaitre

## Abstract

Efferocytosis, the efficient clearance of apoptotic cells (ACs) by phagocytes, is vital for maintaining tissue homeostasis. Here, we reveal the role of the secreted protein NimB1 in reducing apoptotic cell recognition and binding in the early stages of efferocytosis. NimB1 is expressed in macrophages (also called plasmatocytes) and binds to ACs in a phosphatidylserine-dependent manner. Structural analysis shows that NimB1 shares striking similarities with the bridging molecule NimB4, and possesses two phosphatidylserine-binding motifs, supporting its role in efferocytosis. Larval macrophages of NimB1 null mutants display a hyper-phagocytic phenotype characterized by increased engulfment of ACs. Confocal imaging reveals that NimB1 specifically inhibits early steps in internalization of ACs, but does not impact phagosome maturation. We find that NimB1 is a secreted factor that negatively regulates efferocytosis, antagonizing the role of NimB4. Our study and the analogous opposing roles of Draper Isoforms II and I in efferocytosis suggest that a balance of negative and positive regulators allows optimization of the rate of apoptotic cell clearance by macrophages.

## Introduction

Apoptosis and apoptotic cell clearance (efferocytosis) are critical biological processes that maintain cellular and systemic health across multicellular organisms. Apoptosis, the programmed cell death mechanism, plays an essential role in eliminating surplus and damaged cells. Subsequent to apoptosis, efferocytosis facilitates the swift and efficient removal of cellular debris. This process involves the engulfment and degradation of apoptotic corpses by phagocytes, which is crucial for averting inflammatory responses and maintaining tissue homeostasis. Efferocytosis is not only vital for developmental processes but also plays a significant role in immune regulation and tissue homeostasis by preventing the accumulation of cellular waste (Barth et al. 2017; Lemke 2019; Arandjelovic and Ravichandran 2015).

Efferocytosis involves highly complex interactions between ligands and receptors, where phagocytic receptors identify “eat-me” signals on the plasma membranes of dying cells (Elliott and Ravichandran 2016; Boada-Romero et al. 2020; Cockram et al. 2021). Phosphatidylserine (PS), a negatively charged phospholipid typically located in the inner leaflet of the lipid bilayer in healthy cells, is one of the best-characterized “eat-me” signals. In homeostatic conditions, PS is transferred from the outer to the inner leaflet by the flippase ATP11C, maintaining membrane asymmetry. During apoptosis, the inactivation of flippases by caspase-mediated cleavage and the activation of scramblases by similar cleavage events promote relocalization of PS to the outer leaflet, resulting in the high PS exposure characteristic of apoptotic cells (ACs) (Fadok et al. 1998; Segawa and Nagata 2015; Segawa et al. 2014). Binding of phagocytic receptors to PS on an apoptotic cell triggers the formation of a phagocytic cup, initiating uptake and degradation of the ACs. (Callahan, Williamson, and Schlegel 2000). Phagocytes may bind target particles directly through a variety of transmembrane receptors or indirectly by engaging bridging molecules. Bridging molecules are secreted proteins that recognize “eat-me” signals and facilitate the interaction between ACs and the phagocyte’s transmembrane receptors. For instance, the glycoprotein MFG-E8, which is secreted by phagocytic cells, links the integrin receptor αvβ3 to PS, optimizing apoptotic cell uptake (Hanayama et al. 2002; Fuller and Van Eldik 2008; Akakura et al. 2004; Borisenko et al. 2004; Kusunoki et al. 2012). In mice, the absence of MFG-E8 leads to a lupus-like disease due to impaired apoptotic cell clearance (Yamaguchi et al. 2010).

In recent decades, studies of phagocytosis have expanded to genetically tractable model organisms, such as the fruit fly *Drosophila melanogaster* (Brown 2024; Davidson and Wood 2020; Melcarne, Lemaitre, and Kurant 2019; Ulvila, Vanha-Aho, and Rämet 2011). The *Drosophila* genome fully sequenced and extensively annotated, has relatively low redundancy, and is amenable to powerful genetic and molecular techniques as well as high resolution *in vivo* and *ex vivo* imaging. Studies of phagocytosis have highlighted a high degree of similarity between *Drosophila* and mammals, revealing that the process is highly conserved (Melcarne, Lemaitre, and Kurant 2019; Ulvila, Vanha-Aho, and Rämet 2011). Many *Drosophila* phagocytic receptors belong to the Nimrod family, a group of twelve proteins that contain specialized adhesive EGF repeats (NIM repeats), known for their roles in phagocytosis and adhesion (Bork et al. 1996; Somogyi et al. 2008). These Nimrod proteins are clustered mainly on the second chromosome, and are categorized into A-type, C-type and B-type. A-type Nimrods, such as Draper and NimA, and C-type Nimrods, such as NimC1-4 and Eater, are transmembrane proteins implicated in phagocytosis of pathogens and ACs, and in regulation of the adhesion and mobility of plasmatocytes, the *Drosophila* equivalent of macrophages (Kocks et al. 2005; Kurucz et al. 2007; Bretscher et al. 2015; Kurant et al. 2008; Elliott and Ravichandran 2008). Among these transmembrane Nimrods, Eater and NimC1 play a key role in bacterial phagocytosis (Kocks et al. 2005; Kurucz et al. 2007; Bretscher et al. 2015), while NimC4 (also known as SIMU) and Draper (a conserved member of the CED1/MEGF-10 family) bind PS (Kurant et al. 2008; MacDonald et al. 2006; Manaka et al. 2004; 2004; Shklyar et al. 2013; Tung et al. 2013; Mangahas and Zhou 2005; Hamon et al. 2006; Freeman et al. 2003; Fullard, Kale, and Baker 2009). Other *Drosophila* receptors including the integrins βν and αPS3,the CD36 receptor Santa-Maria (Hilu-Dadia et al. 2025), and the bridging molecule Orion (Ji et al. 2023; Nagaosa et al. 2011;) also have demonstrated roles in phagocytosis. The five secreted Nimrod B-type proteins (NimB1-5) encoded by the *Drosophila* genome are less well characterized. Of these, NimB5 is a metabolic factor secreted from the fat body that acts to down-regulate hemocyte proliferation and adhesion upon starvation. Thus, NimB5 tailors resource allocation according to the metabolic state of an animal (Ramond et al. 2020). In contrast, NimB4 has been studied for its role in efferocytosis, and mutants of *NimB4* share phenotypes with *draper* mutants (Manaka et al. 2004; Petrignani et al. 2021; Logan et al. 2012). NimB4 directly binds ACs through PS recognition, serving as a molecular bridge that enhances phagocyte interactions with apoptotic corpses. NimB4 is crucial for both initiation of phagocytosis and for the subsequent phagosome maturation processes (Petrignani et al. 2021). Little is known about the function of NimB proteins other than NimB4 and NimB5; however, the NIM repeats shared by this protein family suggest that they also have roles in macrophages, and specifically in phagocytosis.

Here, we characterize the function of NimB1 and reveal its role in the phagocytosis of ACs.

## Results

### NimB1 shares structural similarities with NimB4

Phylogenetic analysis shows that among the NimB proteins, NimB1 and NimB4 are highly similar and stem from a duplication event, while NimB2 represents an earlier evolutionary split (Kurucz et al. 2007). Both proteins contain seven NIM repeats and a signal peptide, but NimB4 is O-glycosylated while NimB1 is not (Somogyi et al. 2008). Consistent with the phylogenetic analysis, AlphaFold2 revealed that NimB1 and NimB4 exhibit striking structural similarities not shared by NimB2 (**Figure 1 A-B**). The sequence of NimB1 aligns over 344 residues with NimB4 (∼85% of its sequence), compared to only 165 residues (∼41%) for NimB2. The low RMSD (Root Mean Square Deviation) value (2.70 Å for NimB1 vs. NimB4) and high TM-score (Template Modelling score) of 0.761 indicate a strong structural resemblance, supporting the notion that these proteins may share a close functional relationship (Zhang and Skolnick 2005). These metrics stand in contrast to the weaker similarity observed between NimB1 and NimB2 (TM-score = 0.322, RMSD = 4.59 Å) (**Figure 1 A-B**). Interestingly, both NimB1 and NimB4 contain an RRY motif previously shown to mediate PS binding in the bridging molecule Orion (Ji et al. 2023). NimB1 contains two such motifs, while NimB4 has one (**Figure 1 C-D**). Phylogenetic and structural similarities between NimB1 and NimB4 suggest that NimB1 may serve a function in apoptotic cell clearance similar to NimB4.

**Figure 1:**
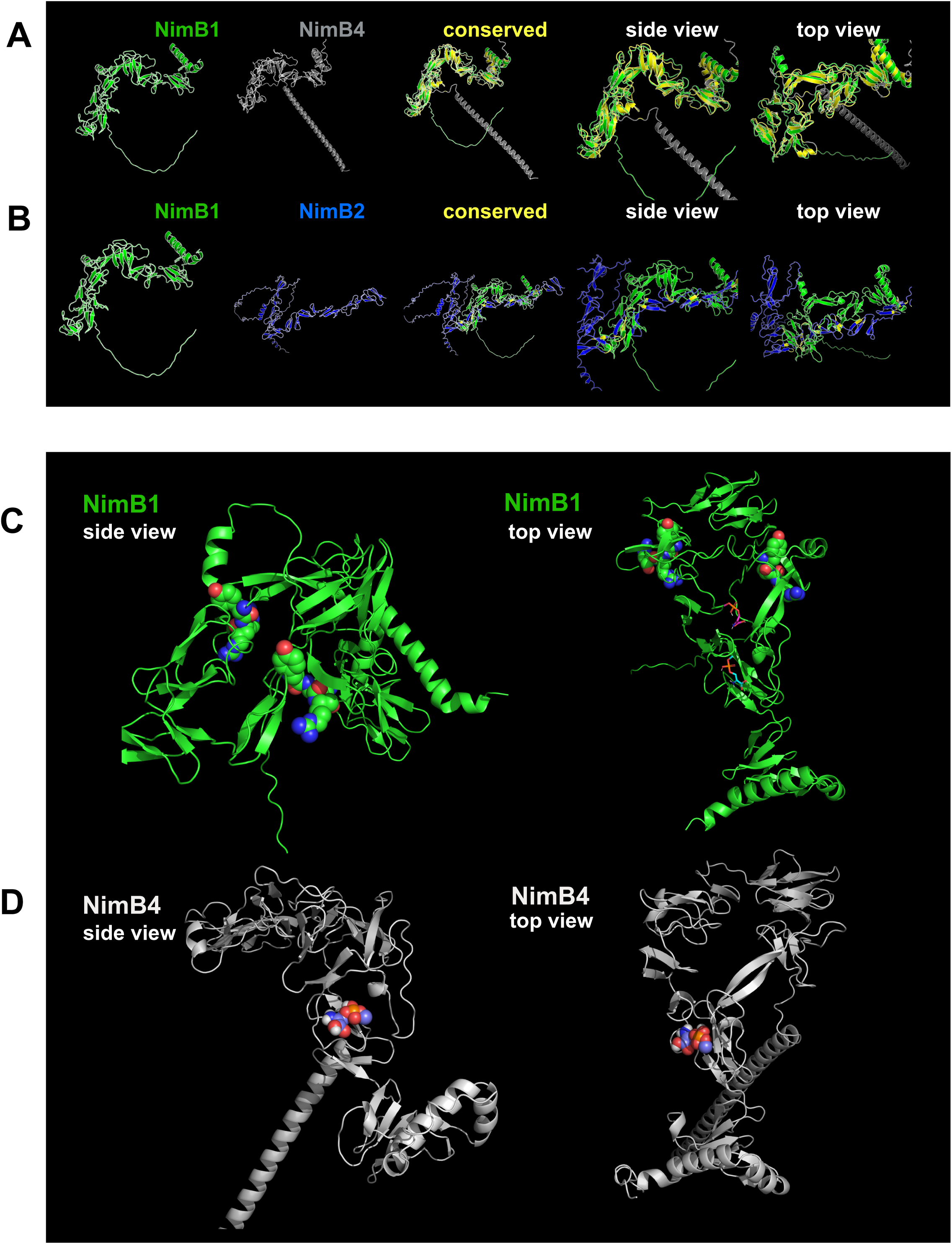
Structural alignment and PS-binding motif visualization in NimB1 and NimB4. (A, B) Structural alignment of NimB1 and NimB4 (A) or NimB1 and NimB2 (B) modeled using AlphaFold2. Conserved regions between aligned structures are shown in yellow. NimB1 is depicted in green, NimB4 in gray, and NimB2 in blue. Side and top views are provided for each comparison to illustrate the conserved structural features. (C, D) High-resolution views of NimB1 (C, green) and NimB4 (D, gray), showing their side and top orientations. The RRY motif, implicated in PS binding and apoptotic cell clearance (Hui Ji et al., 2023), is visualized in a space-filling representation, while the rest of the protein is depicted in cartoon style. NimB1 contains two RRY motifs, while NimB4 contains one, suggesting functional similarity in PS binding.

### NimB1 is expressed in macrophages in a pattern consistent with a role in efferocytosis

During the *Drosophila* life cycle, significantly higher apoptosis is observed during embryogenesis and pupariation due to tissue remodeling (Shklover, Levy-Adam, and Kurant 2015; Serizier and McCall 2017). Some receptors such as SIMU predominantly operate during embryogenesis, whereas Draper functions across all life stages (Kurant et al. 2008). To analyze the expression dynamics of *NimB1*, RT-qPCR was performed across developmental stages. Similar to *NimB4* and *simu* (Kurant et al. 2008; Petrignani et al. 2021), *NimB1* expression peaks during embryogenesis (**Figure 2A**). In third instar larvae, tissue-specific analysis revealed that *NimB1* is predominantly expressed in macrophages, in line with previous findings that these macrophage-like cells are the primary site of *NimB1* and *NimB4* expression (Petrignani et al. 2021; Ramond et al. 2020) (**Figure 2B**). RT-qPCR analysis also revealed that *NimB1* is induced in third instar larvae one hour following clean injury, similar to *NimB4* (Petrignani et al. 2021) (**Figure 2C**).

**Figure 2.**
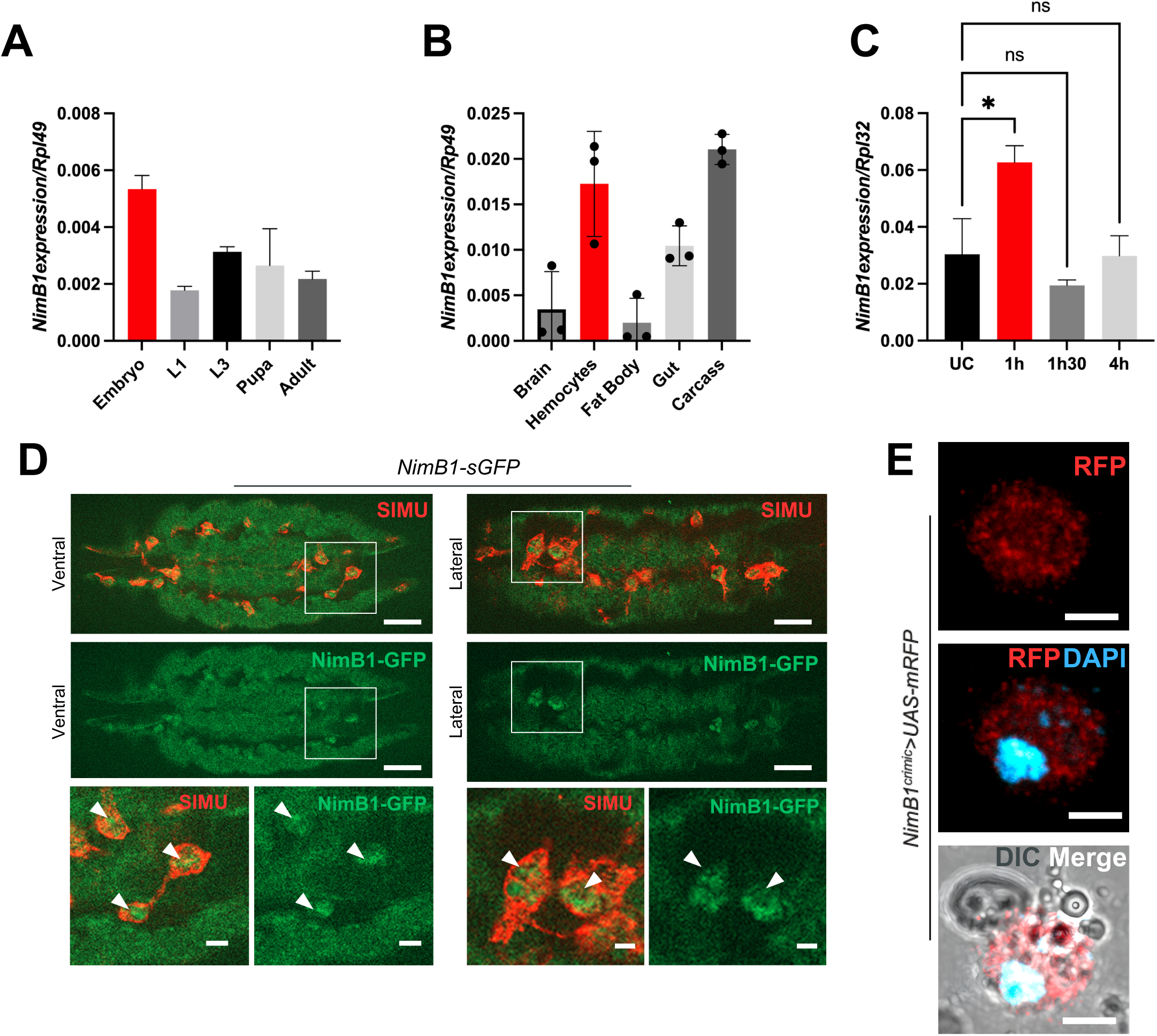
The expression pattern of NimB1 in macrophages is consistent with a role in efferocytosis. (A) RT-qPCR analysis of *NimB1* transcripts, normalized to *Rp49* from embryo, instar larvae 1 (L1), instar larvae 3 (L3), pupa and adult wild-type *Drosophila*. Data are represented as mean ± SD from three independent experiments. (B) RT-qPCR analysis of *NimB1* transcripts, normalized to *Rp49*, extracted from brain, macrophages, fat body, gut and carcass of wild-type third instar wandering larvae. Data are represented as mean ± SD from three independent experiments. (\**P* < 0.05, ns = not significant, by one-way ANOVA with Tukey’s post hoc test) (C) Quantification of *NimB1* transcript expression relative to *Rpl32* in macrophages, in unchallenged conditions (UC) and at time points following clean injury. (D) Representative confocal imaging of *NimB1-sGFP* macrophages from stage 16 embryos, ventral and lateral views. Tissues were stained with anti-GFP (green, corresponding to *NimB1-sGFP*) and anti-SIMU antibodies (red, corresponding to macrophages). The arrowheads show the presence of NimB1-sGFP inside the embryonic macrophages. (E) Live confocal imaging of macrophages of *NimB1^crimic^* flies overexpressing *UAS-mRFP* under the control of the *TI{CRIMIC.TG4.2}NimB1^CR02514-TG4.2^*. Scale bar = 10 μm.

To further confirm the expression pattern of *NimB1*, we generated a transgenic *Drosophila* line carrying a V5-sGFP-tagged *NimB1* fusion under its own regulatory sequences, derived from the Dresden pFlyfos collection (Sarov et al. 2016). Confocal imaging demonstrated that NimB1-sGFP localized in macrophages during embryogenesis (**Figure 2D**), similar to NimB4 (Petrignani et al. 2021). However, GFP signals were undetectable in third instar larvae, likely due to low *NimB1* expression at this stage. Instead, we used a *NimB1^crimic^* fly line, where the first intron of the *NimB1* gene bears a CRIMIC cassette insertion containing a GAL4 reporter (P.-T. Lee et al. 2018). Macrophages dissected from *NimB1^crimic^* larvae overexpressing mRFP [;*NimB1^crimic^/+;UAS-mRFP1/+]* confirmed that *NimB1* is expressed at this stage (**Figure 2E**).

Overall, our study reveals that *NimB1* is expressed in macrophages with peak expression during embryogenesis, suggesting a role in tissue remodeling. The expression pattern of NimB1 is similar to NimB4, where both are induced upon clean injury.

### NimB1 is a secreted protein

To explore the subcellular localization of NimB1 we overexpressed a *UAS-NimB1-RFP* construct using the hemocyte driver *Hml^p2a^-Gal4* (Stephenson et al. 2022). The *UAS-SP^vkg^-RFP* line in which RFP is fused to the signal peptide of Viking, a subunit of Collagen IV protein (Ke et al. 2018), was used as a control. The NimB1-RFP signal exhibited a vesicular localization in third instar larval macrophages **(Supplementary document S1 A-D)**, and was also quantitatively enriched at plasma membrane regions **(Supplementary document S1 B-E)**, suggesting that NimB1 is secreted. Furthermore, a strong NimB1-RFP signal was detected in nephrocytes of larvae expressing the *UAS-NimB1-RFP* construct from the fat body using the *Lpp-GAL4* driver **(Supplementary document S1 F)**. As nephrocytes are known to filter hemolymph proteins (Troha et al. 2019; Ivy et al. 2015), this confirms that NimB1-RFP was secreted to the hemolymph when expressed in the fat body.

In sum, the vesicular localization of NimB1 and its enrichment at the plasma membrane and in the extracellular space confirms that consistent with its signal peptide, NimB1 is processed and transported for secretion into the extracellular environment.

### NimB1 directly binds to ACs through phosphatidylserine

Phagocytic receptors often have a low affinity for the target ligand (Koopman et al. 1994). To compensate for this low affinity, binding of ACs typically involves the sequential engagement of multiple low-affinity receptors and is facilitated by bridging molecules (Uribe-Querol and Rosales 2020; Borisenko et al. 2004; Akakura et al. 2004; Fuller and Van Eldik 2008; Tollis et al. 2010; Yamaguchi et al. 2010; Petrignani et al. 2021; Ji et al. 2023). The presence of PS binding pocket in NimB1 prompted us to test its ability to bind ACs. Hemolymph from third instar larvae expressing *UAS-NimB1-RFP* under the *Hml^p2a^-Gal4* driver was co-incubated with Carboxyfluorescein Succinimidyl Ester (CFSE)-labeled ACs. Confocal imaging and 3D reconstructions revealed that NimB1 binds ACs in a distinct patch-like pattern, covering the cell surface (**Figure 3A-C**).

**Figure 3.**
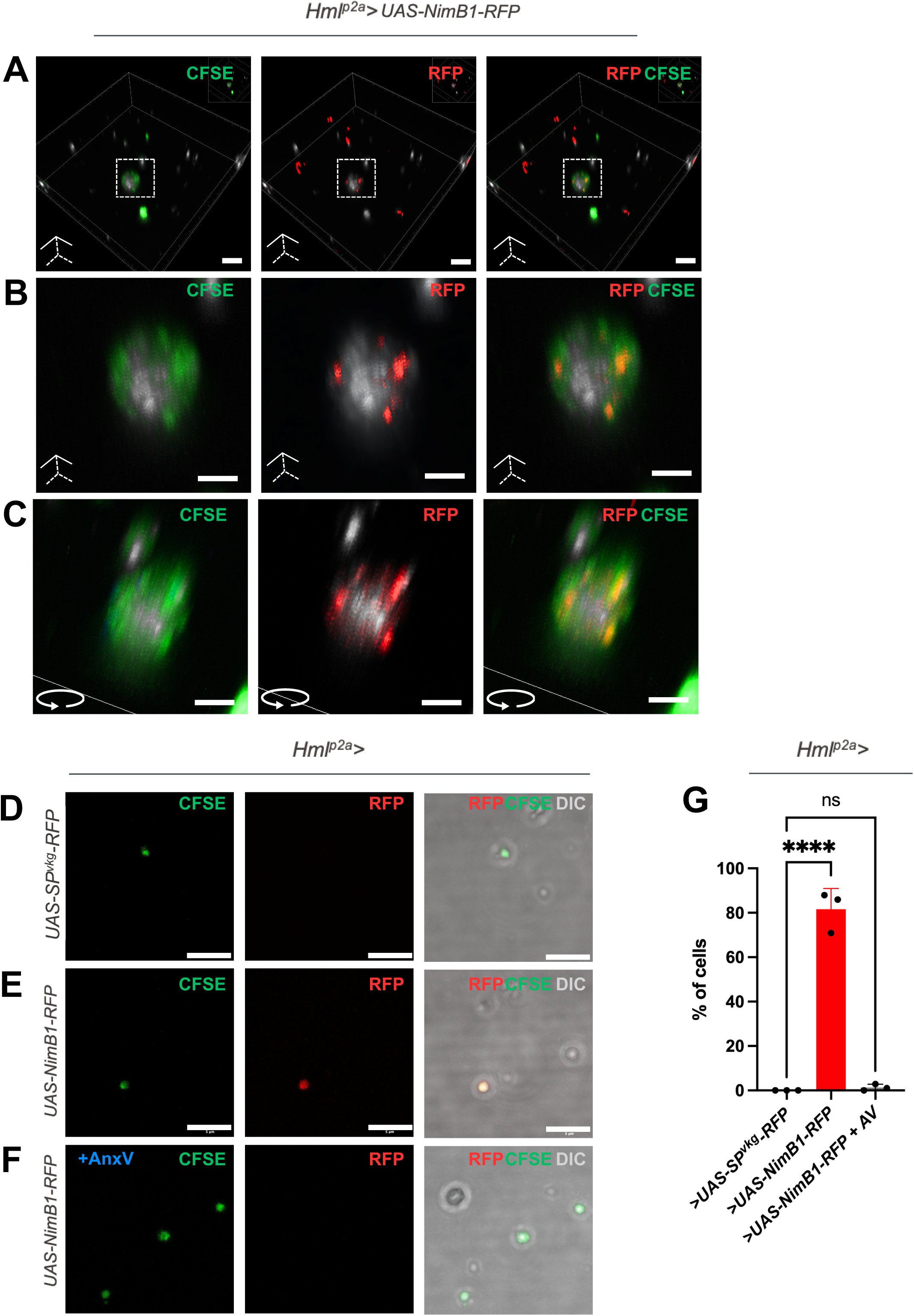
NimB1 binds to ACs in a PS-dependent manner. (A-C) Representative confocal microscopy images of fixed macrophages expressing *UAS-NimB1-RFP* under the control of the hemocyte *Hml^p2a^-Gal4* driver, stained with an anti-RFP antibody and co-incubated with ACs stained with CFSE. Scale bar = 5 μm. Overlay of fluorescence and differential interference contrast (DIC), 3D rendering completed with Imaris Viewer. (B-C) Zoomed image of the area. Scale bar = 3μm. (C) Angled zoom image. (D-F) Representative confocal images of CFSE-stained apoptotic bodies (green) incubated with secreted *UASNimB1-RFP* (E,F) or *SP^vkg^-RFP* (D) from macrophages with *Hml^p2a^-Gal4* driver. Apoptotic bodies were pre-incubated in absence (D,E) or presence (F) of Annexin V (25 μg/ml). Scale bar = 5 μm. (G) Quantification of the colocalization of NimB1-RFP or SP^vkg^-RFP with apoptotic bodies in the presence or absence of Annexin V, as measured by the percentage of AC colocalized with the RFP signal. Values from three independent experiments are represented as mean ± SD. (\*\*\*\**P* < 0.0001, ns = not significant, by one-way ANOVA with Tukey’s post hoc test).

To determine whether NimB1 binding to ACs is PS-dependent, we pre-incubated in the same experimental settings ACs with Annexin V, a protein with high PS affinity (van Engeland et al. 1998; Janko et al. 2013). Annexin V significantly reduced NimB1 binding, confirming that NimB1 interacts with ACs through PS (**Figure 3G**). The specificity of NimB1 binding is further validated by control experiments showing that the secreted protein *UAS-SP^vkg^-RFP*, which does not have affinity to PS, did not similarly bind apoptotic corpses (**Figure 3D-G**). This experiment demonstrates that NimB1 binds ACs in a PS-dependent manner, at a lower affinity than Annexin V.

### NimB1-deficient macrophages display increased adhesive properties

To analyze the *in vivo* functions of NimB1 in efferocytosis, we generated a *NimB1^229^* null mutant using CRISPR-Cas9 technology, in addition to the previously mentioned *NimB1^crimic^* line which functions as a null mutant due to the CRIMIC cassette insertion in the first exon of the gene. *NimB1^229^* has a deletion of 11 base pairs in the first exon, resulting in a frameshift leading to a premature stop codon. *NimB1^229^* and *NimB1^crimic^* flies were backcrossed into the *w^iso^* (DrosDel) background. *NimB1^229^* and *NimB1^crimic^* homozygous mutants were lethal at the second larval instar and pupal stage, respectively. However, transheterozygous mutants [*NimB1^229^*/*NimB1^crimic^*] were viable until adulthood, providing an opportunity to explore the functions of NimB1 in *NimB1^crimic^*homozygous larvae and in *NimB1^229/crimic^* larvae and adults. The independent lethality of the two *NimB1* mutant lines may be due to complex genetic interaction in the *Nimrod* genomic clusters.

Given the similarities between NimB1 and NimB4, we also generated a *NimB1, NimB4* double mutant by injecting a gRNA targeting the *NimB1* sequence in *NimB4^sk2^* embryos, using the CRISPR-Cas9 technique. The *NimB1^61^, NimB4^sk2^*double mutant has deletions in the first exons of both *NimB1* and *NimB4* **(Supplementary document S2A)**, and this line was viable until adulthood.

As NimB5 is implicated in hemocyte proliferation (Ramond et al. 2020), we quantified hemocyte number in *NimB1* and *NimB1, NimB4* double mutants using Fluorescence-Activated Cell Sorting (FACS) analysis. Third instar larval macrophages from wild-type, *NimB1^crimic^*, *NimB1^229/crimic^*and*, NimB4^sk2^* mutants display hemocyte count comparable to wild-type controls, suggesting that similar to NimB4 (Petrignani et al. 2021), NimB1 does not play a role in hematopoiesis **(Supplementary document S3 A)**. However, *NimB1, NimB4* double mutant larvae showed a significant increase in hemocyte number, indicating that these genes may have a cryptic direct or indirect effect on hematopoiesis.

To further investigate the role of NimB1, we examined its impact on hemocyte adhesion and cytoskeletal organization. Macrophages from wild-type, *NimB1* and *NimB4,* single and double mutants were seeded onto glass, allowed to adhere, and stained with phalloidin to visualize actin structures. Macrophages derived from *NimB1^crimic^*and *NimB4^sk2^* single mutant or *NimB1^61^, NimB4^sk2^* double mutants exhibited significantly increased spreading compared to wild-type cells **(Supplementary document S3 B, D),** indicating enhanced adhesive properties. Analysis of actin structures revealed an increased ratio of filopodia to lamellipodia across all mutants **(Supplementary document S3 C).** This shift in cytoskeletal dynamics suggests enhanced filopodial projection formation, reflecting a reorganization of the actin cytoskeleton. These changes may enhance the ability of mutant macrophages to adhere to surfaces and interact with target particles.

### NimB1 delays the early-stage efferocytosis

Given the structural and evolutionary similarities between NimB1 and NimB4, we hypothesized that NimB1 might also contribute to efferocytosis. To test this, we performed an *ex vivo* phagocytic assay using macrophages from wild-type, *NimB1^crimic^, NimB1^229^*^/*crimic*^*, NimB4^sk2^*,*NimB1^61^, NimB4^sk2^* and *draper*^Δ*5*^ larvae. CFSE-labeled ACs were incubated with macrophages bled from third instar larvae for 1 or 2 hours, and phagocytic index was calculated by multiplying the percentage of macrophages with internalized ACs by their mean fluorescence intensity (MFI) **(Supplementary document S4).**

Surprisingly, *NimB1* mutants displayed a hyper-phagocytic phenotype, with significantly increased MFI across all time points (**Figure 4 A, B**). As expected (Freeman et al. 2003; Petrignani et al. 2021), *draper* mutants displayed impaired apoptotic cell binding and engulfment, while *NimB4^sk2^* mutants showed a reduction in phagocytic activity at later time points that was not statistically significant in our hands (**Figure 4D**). To confirm the hyper-phagocytic phenotype of *NimB1* mutant macrophages, we performed rescue experiments by reintroducing *NimB1* or *NimB1-RFP* under the control of the *NimB1* promoter in the *NimB1^crimic^* mutant background [*NimBc^crimic^ (Gal4)*; *UAS-NimB1*]. Both constructs successfully restored phagocytic activity to wild-type levels, as evidenced by both MFI and phagocytic index (**Figures 4C and 4D, Supplementary document S5**). This rescue reinforces the notion that NimB1 inhibits phagocytosis of ACs, opposing the role of NimB4.

**Figure 4.**
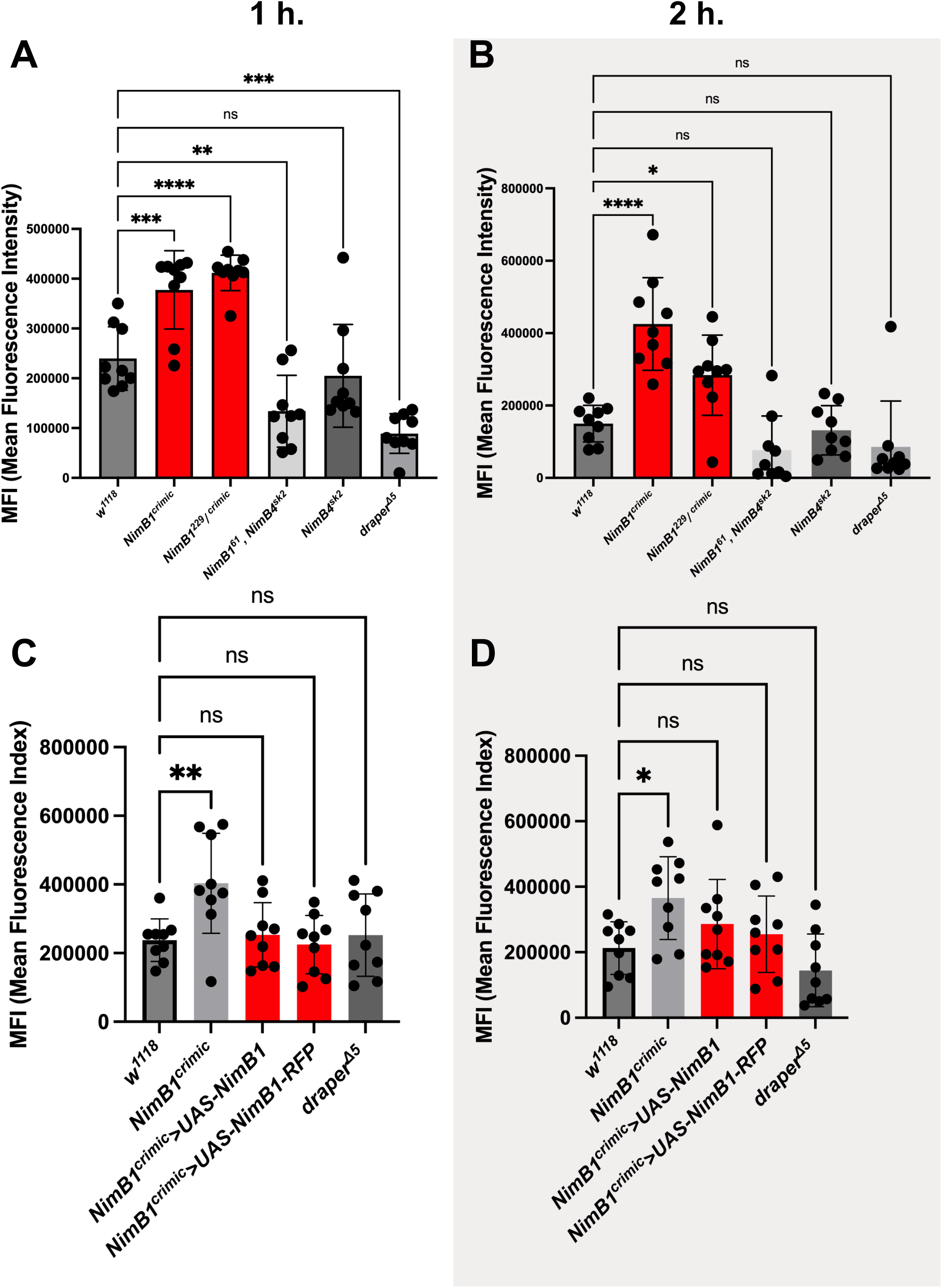
Enhanced phagocytic activity in *NimB1* mutants during apoptotic cell uptake. (A, B) Mean Fluorescence Intensity (MFI) of CFSE-labeled ACs internalized by macrophages from *w^1118^* (wild-type), *NimB1^crimic^*, *NimB1^Trans^*, *NimB4^sk2^*, and *draper*^Δ*5*^ larvae after 1 hour (A) and 2 hours (B) of incubation at room temperature, measured using flow cytometry. Data are presented as mean ± SD from three independent experiments. (\*\*\*\**P* < 0.0001, \*\*\**P* < 0.001, \*\**P* < 0.01, \**P* < 0.05, ns = not significant, by one-way ANOVA with Tukey’s post hoc test). (C, D) Rescue experiments in *NimB1^crimic^* macrophages expressing *UAS-NimB1-RFP* or *UAS-NimB1* under the control of the *NimB1* promoter in the mutant *NimB1^crimic^*background. Macrophages and CFSE-labelled ACs were incubated for 1 hour (С) or 2 hours (D). (\*\**P* < 0.01, \**P* < 0.05, ns = not significant, by one-way ANOVA with Tukey’s post hoc test).

To explore the dynamics of early apoptotic cell clearance, we repeated the *ex vivo* phagocytosis assay at 10, 15, and 30-minute time points. Three stages of phagocytic events were quantified: attachment, where the ACs are in contact with but not fused to the plasma membrane of the phagocyte; membrane binding, where the bound apoptotic cell is no longer a distinct entity due to membrane fusion; and full internalization, where the apoptotic cell signal is completely inside the cytosol of the phagocyte (**Figure 5A**). Strikingly, macrophages from *NimB1* mutants progressed more rapidly to internalization, with a significant increase in numbers of fully internalized ACs as early as 10 minutes post-incubation (**Figure 5B-D**). This suggests that wild-type NimB1 delays the transition from recognition to engulfment, consistent with the high internalization rate found in *NimB1* mutant macrophages at the 1 and 2 hour time points. In contrast, *NimB4* deficient macrophages exhibited delayed internalization, with minimal internalization of ACs at early time points increasing to a near wild-type level by 30 minutes (**Figure 5B-D**). Double mutants (*NimB1^61^, NimB4^sk2^*) mirrored the *NimB4* phenotype, showing delayed attachment and internalization. As expected, *draper* mutant larval macrophages displayed minimal apoptotic cell uptake across all time points (**Figure 5B-D**). Quantification of these events revealed that NimB1-deficient macrophages exhibited similar levels of apoptotic cell attachment compared to wild-type, but displayed a pronounced increase in internalization efficiency.

**Figure 5.**
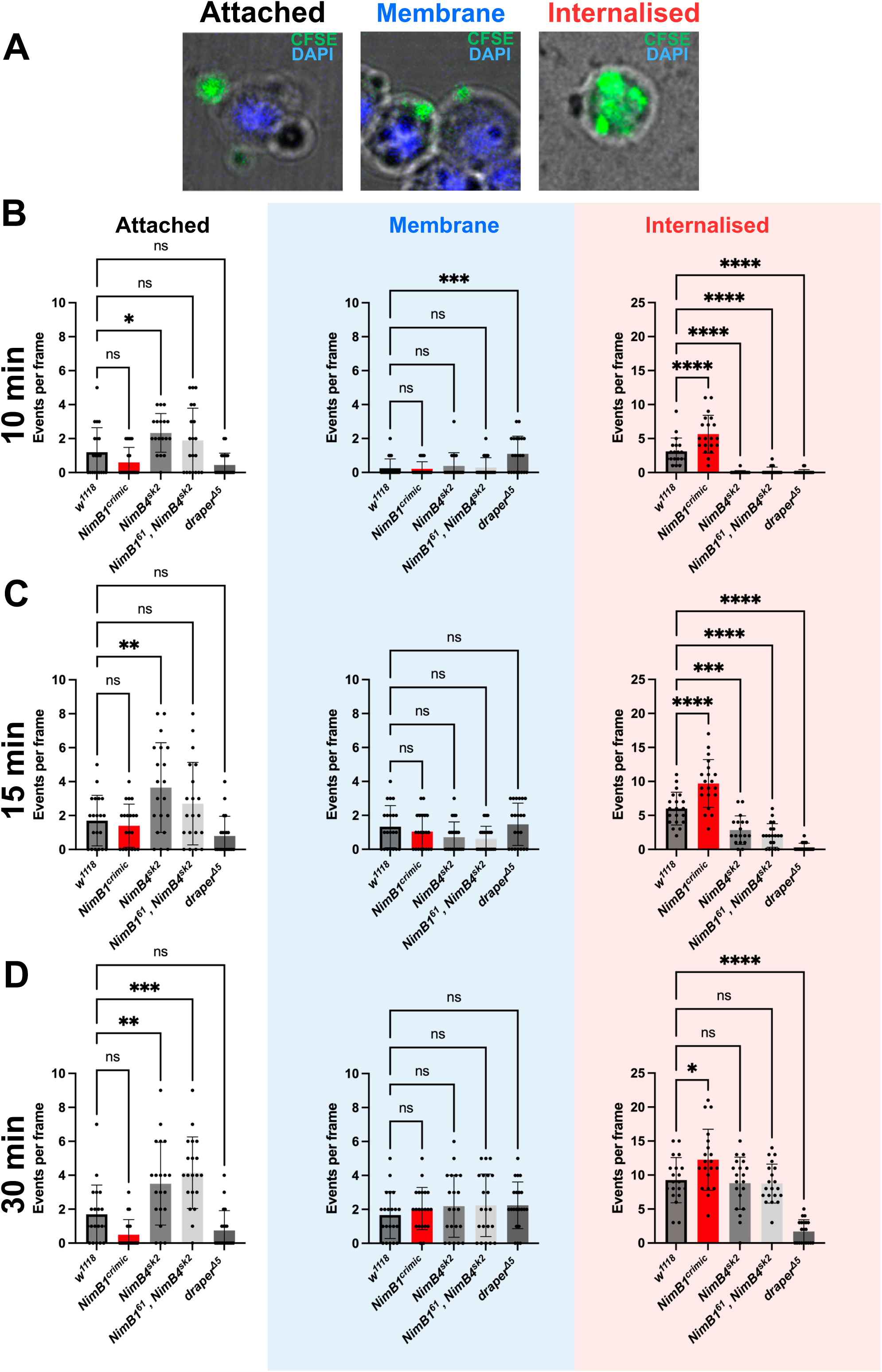
Rapid internalization of apoptotic cells in *NimB1* mutant macrophages at early phagocytosis stages. (A) Representative confocal images of macrophages dissected from third instar larvae and incubated with CFSE-labeled ACs for 10, 15, and 30 minutes. Interactions of ACs and macrophages were classified into three categories: attached (gray), at the plasma membrane (blue), and fully internalized (red) into macrophages. Green: AC CFSE staining; Blue: DAPI staining. (B-D) Quantification of the number of phagocytic events of ACs per frame, categorized as attached, membrane-associated, or internalized, for each genotype across three time points: (B) 10 minutes, (C) 15 minutes, and (D) 30 minutes. Data are derived from three independent experiments and represented as individual data points with means ± SD. (\**P*<0.05, ** *P* <0.01, *** *P* <0.001, **** *P* <0.0001, ns = not significant, by one-way ANOVA with Tukey’s post hoc test).

We then investigated the role of NimB1 in apoptotic cell clearance under *in vivo* conditions, by performing a live phagocytosis assay following mechanical injury. Third instar larvae were pinched with forceps, a procedure that induces apoptosis and increases the availability of apoptotic material in the hemolymph. Macrophages were extracted prior to challenge and at 90 minutes post-injury, and internalized apoptotic cell fragments were identified by the presence of DAPI signals in the cytoplasm of the macrophages, distinct from the cell’s nucleus (**Figure 6A, B**). Under unchallenged conditions, wild-type macrophages exhibited minimal accumulation of apoptotic cell fragments (**Figure 6C**). In contrast, *NimB4^sk2^, NimB1^61^,NimB4^sk2^*, and *draper*^Δ*5*^ mutants displayed significant accumulation of apoptotic cell fragments even in the absence of apoptotic challenge, indicating a defective baseline clearance of apoptotic material.

**Figure 6.**
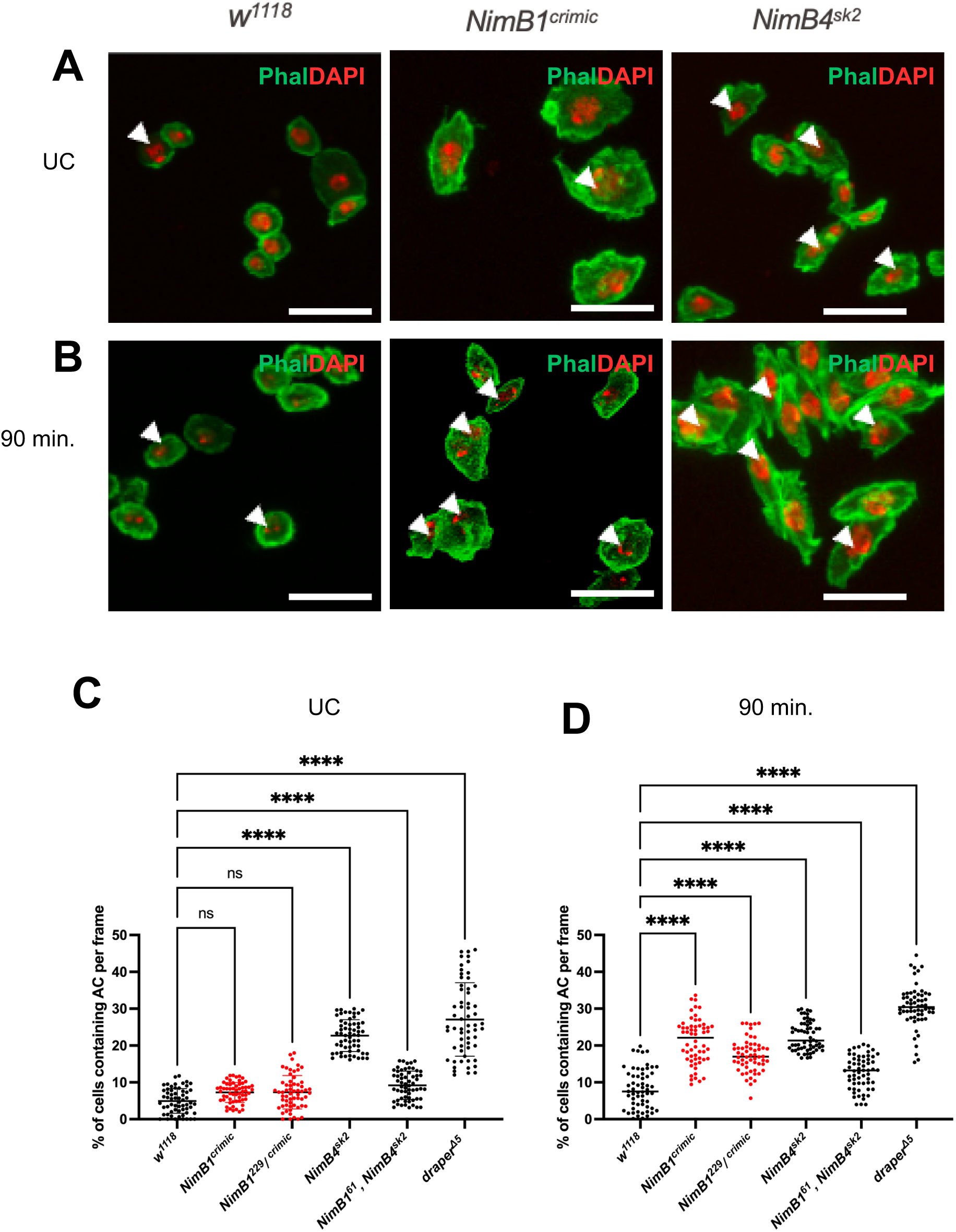
*In vivo* increased apoptotic cell uptake by *NimB1* mutant macrophages upon pinching of larvae. (A, B) Representative confocal microscopy images of macrophages isolated from *w^1118^*, *NimB1^crimic^,* and *NimB4^sk2^* larvae. Macrophages were fixed and stained with Phalloidin (green) to label F-actin and DAPI (red) to detect AC fragments. Arrowheads highlight AC fragments. (A) Macrophages in unchallenged (UC) conditions show baseline apoptotic uptake. (B) Macrophages analyzed 90 minutes after larval pinching, showing increased accumulation of DAPI-positive apoptotic fragments in *NimB1* and *NimB4^sk2^* mutants. Scale bars = 10μm. (C, D) Quantification of the percentage of macrophages containing apoptotic corpses under (C) unchallenged (UC) and (D) 90-minute post-pinching conditions. Data are derived from three independent experiments and represented as individual data points with means ± SD. (\*\*\*\**P* < 0.0001, ns = not significant, by one-way ANOVA with Tukey’s post hoc test).

Consistent with the hyper-phagocytic phenotype observed in *ex vivo* assays, macrophages from *NimB1^crimic^* and *NimB1^229/crimic^* mutants accumulated apoptotic fragments at significantly higher levels following mechanical injury compared to controls (**Figure 6D**). Notably, *NimB4^sk2^, NimB1^61^,NimB4^sk2^*, and *draper*^Δ*5*^ mutants failed to internalize additional apoptotic material upon injury, maintaining internalization levels comparable to those seen in unchallenged conditions. These findings confirm that NimB1 plays a unique role in modulating the efficiency of early-stage apoptotic cell uptake, where absence of NimB1 leads to accelerated internalization. In contrast, NimB4 and Draper are required to maintain basal clearance rate and facilitate routine apoptotic cell turnover under both hemostatic and challenged conditions.

### Loss of NimB1 enhances recognition and attachment of ACs

To further confirm the role of NimB1 in the early stages of apoptotic cell clearance, we performed a cold binding assay designed to quantify the ability of macrophages to bind ACs without progressing to internalization. Macrophages isolated from wild-type, *NimB1^crimic^*, and *draper*^Δ*5*^ larvae were incubated with CFSE-labeled ACs at 4°C, a temperature that inhibits active phagocytosis without disrupting binding (Pearson et al. 2003). Interestingly, *NimB1* mutant macrophages exhibited a significant increase in the number of bound ACs compared to wild-type macrophages, suggesting that loss of *NimB1* enhances recognition and attachment (**Figure 7A-B**). By contrast, *draper*^Δ*5*^ mutant macrophages known for defective apoptotic cell recognition displayed minimal binding under the same conditions.

**Figure 7.**
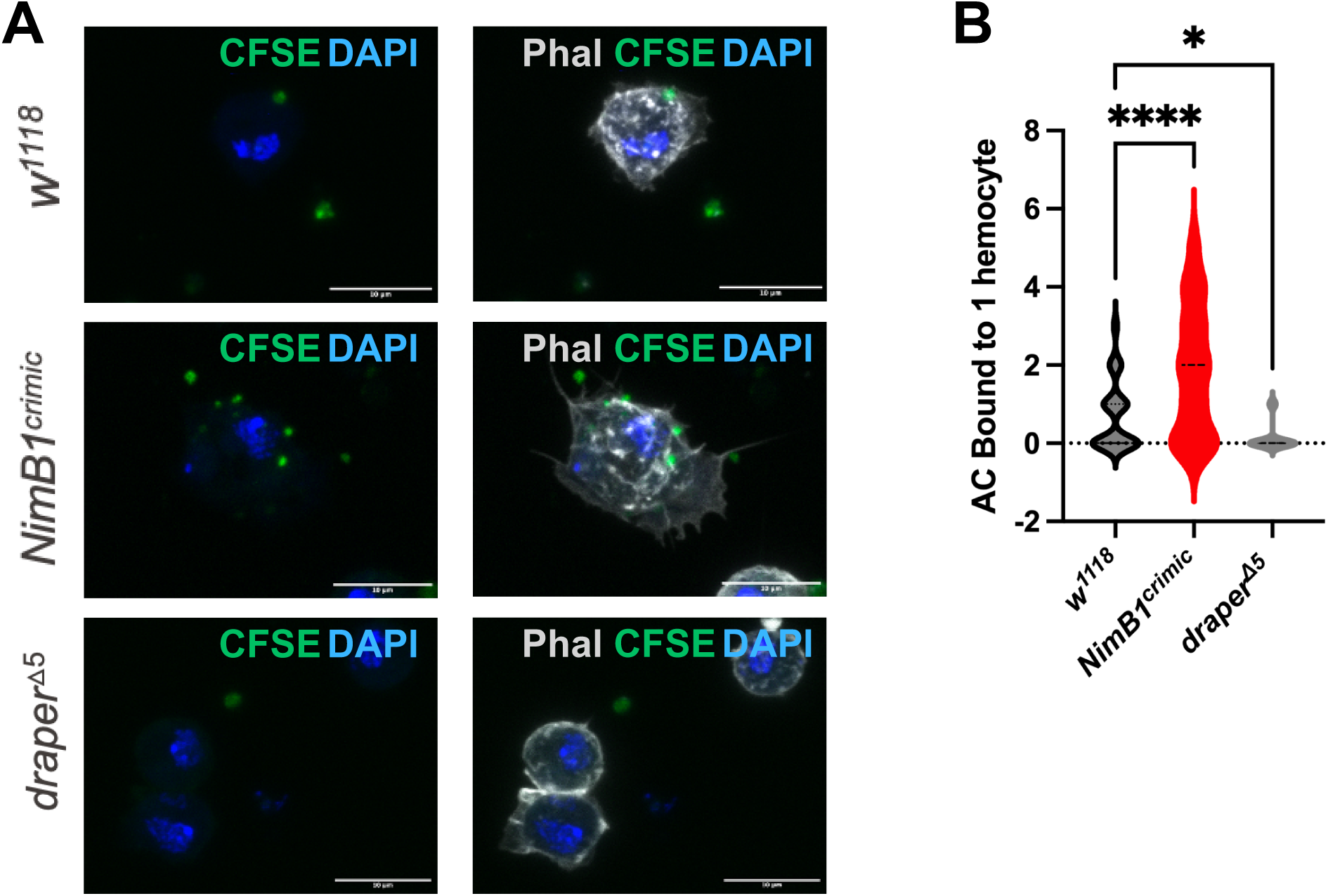
*NimB1* mutant macrophages show enhanced binding to apoptotic corpses. (A) Representative confocal microscopy images of macrophages isolated from *w^1118^*, *NimB1^crimic^*, and *draper*^Δ*5*^ larvae in presence of CFSE-labeled ACs (green) at 4°C. DAPI (blue), phalloidin (gray) to label F-actin. (B) Quantification of ACs bound per hemocyte across genotypes. Data are represented as violin plots, derived from three independent experiments. Dashed lines indicate the median values. (\*\*\*\**P* < 0.0001, \**P* < 0.05, by one-way ANOVA with Tukey’s post hoc test).

Transmission electron microscopy (TEM) revealed that macrophages from *NimB1^crimic^* mutants displayed increased cell surface remodeling compared to wild-type, with prominent phagocytic cup formation (**Figure 8A**). These structures are indicative of enhanced phagocytic readiness and align with a greater capacity for apoptotic cell uptake. In contrast, numerous intracellular vacuoles were found in *NimB4* mutant macrophages, consistent with disrupted phagosome maturation (Petrignani et al. 2021). The *NimB1^61^, NimB4^sk2^*double mutant combined the phenotypes of the single mutants, displaying both enhanced phagocytic cup formation and increased intracellular vacuoles, suggesting distinct yet complementary roles for NimB1 and NimB4 in phagocytosis.

**Figure 8:**
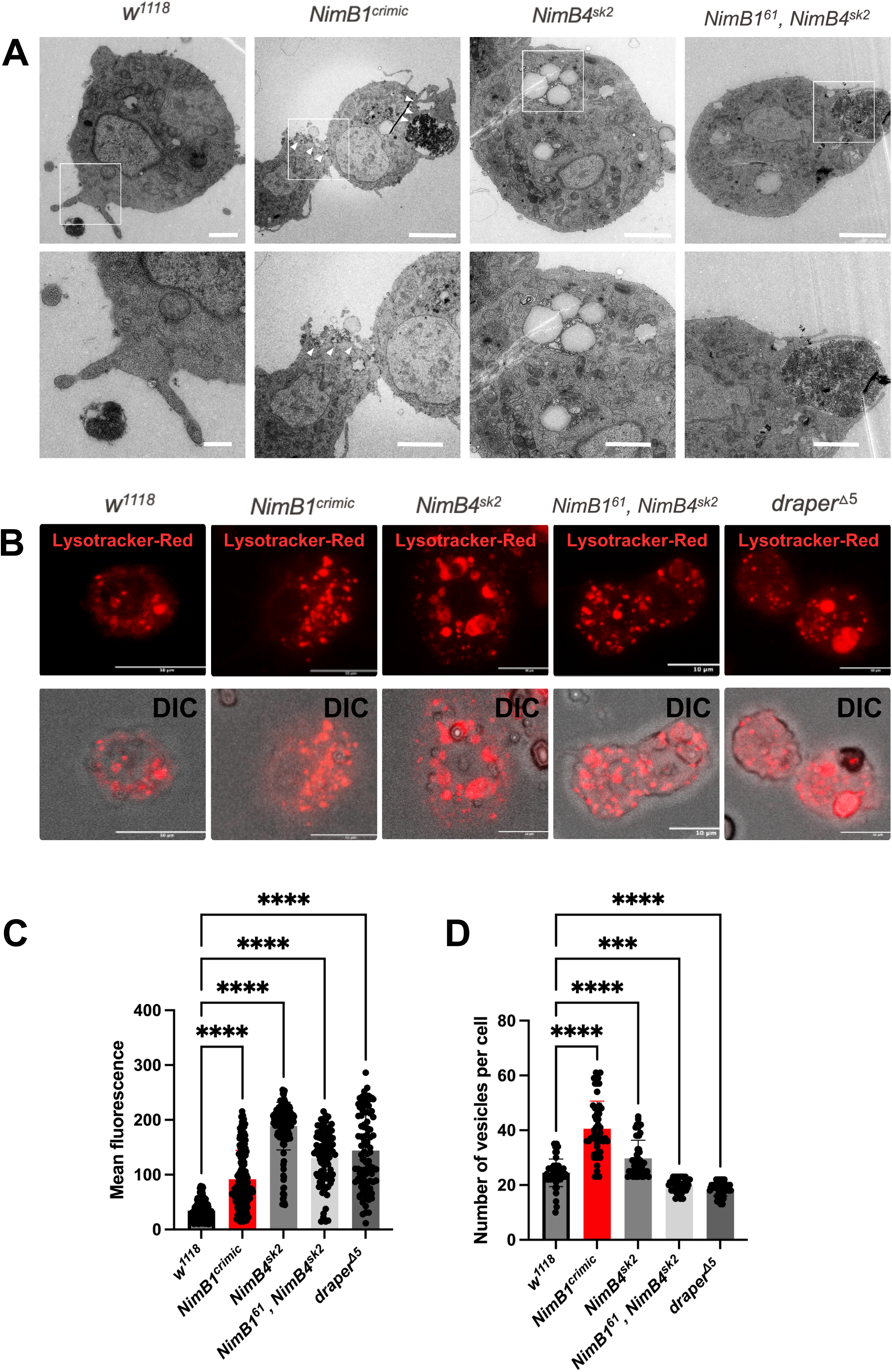
TEM and LysoTracker Red analysis reveal altered phagosome dynamics in *NimB1* and *NimB1^61^,NimB4^sk2^* mutants. (A) Representative TEM images of macrophages from *w^1118^*, *NimB1^crimic^*, *NimB4^sk2^, NimB1^61^,NimB4^sk2^* larvae. Scale bar = 2 μm. Zoomed areas are indicated with dashed rectangles. (B) LysoTracker Red fluorescence (top row) and DIC merged images (bottom row) for *w^1118^*, *NimB1^crimic^*, *NimB4^sk2^, NimB1^61^,NimB4^sk2^* larval macrophages. Scale bar = 5 μm. (C) Quantification of mean LysoTracker Red fluorescence per hemocyte for each genotype. Data are derived from three independent experiments and represented as individual data points with means ± SD. (\*\*\*\**P* < 0.0001, by one-way ANOVA with Tukey’s post hoc test). (D) Quantification of the number of acidic vesicles per hemocyte across genotypes. Data are derived from three independent experiments and represented as individual data points with means ± SD. (\*\*\*\**P* < 0.0001, by one-way ANOVA with Tukey’s post hoc test).

Collectively, our observations reveal that NimB1 delays early recognition and binding of ACs, opposing the phagocytosis-enhancing function of NimB4.

### NimB1 does not contribute to bacterial phagocytosis

Several receptors, notably NimC1, Eater and Draper, have been implicated in the phagocytosis of bacteria (Hashimoto et al. 2009; Bretscher et al. 2015; Melcarne et al. 2019). Having demonstrated a role for NimB1 in apoptotic cell clearance, we next investigated whether NimB1 also contributes to phagocytosis of bacteria. We conducted an *ex vivo* phagocytosis assay using Alexa488-labeled *Escherichia coli* (Gram-negative) and *Staphylococcus aureus* (Gram-positive) particles. As previously described (Bretscher et al. 2015; Melcarne et al. 2019), macrophages from third instar larvae were co-incubated with bacterial particles for 1 hour, and the phagocytic index was calculated. Bacterial phagocytosis was as wild-type in *NimB1^crimic^* mutant larval macrophages, indicating no role for NimB1 in bacterial clearance **(Supplementary document S6)**. *NimB1^61^, NimB4^sk2^* double mutants were also behaving as wild-type, agreeing with prior findings that NimB4 does not contribute to bacterial phagocytosis (Petrignani et al. 2021). In contrast, *NimC1; eater^1^* double mutants exhibited a marked reduction in bacterial uptake, consistent with their established roles as receptors in bacterial phagocytosis.(Kocks et al. 2005; Bretscher et al. 2015; Melcarne et al. 2019)

### NimB1 Increases phagosome formation without affecting maturation

We next investigated the impact of NimB1 on downstream steps of the phagocytic process by examining its role in phagolysosome formation. To assess this, macrophages were stained with LysoTracker Red, a fluorochrome that labels acidic vesicles, including phagosomes and lysosomes (Kocks et al. 2005; Bretscher et al. 2015; Melcarne et al. 2019). In *NimB1^crimic^*mutant macrophages, we observed an elevated number of LysoTracker-positive vesicles with slightly increased fluorescence intensity compared to wild-type controls (**Figure 8B–D**). In contrast, *NimB4*^sk2^ and *draper^Δ5^* mutants displayed enlarged and highly fluorescent phagosomes (**Figure 8B-D**), consistent with a blockage at the phagosome maturation step (Kurant et al. 2008; Petrignani et al. 2021).

These results suggest that the increased LysoTracker signal in *NimB1* mutants reflects enhanced phagocytic activity, specifically an increase in the number of phagosomes formed, rather than a defect in phagosome maturation. The acidic vesicles in *NimB1* mutants were wild-type in size and fluorescence intensity, indicating that the maturation process proceeds normally. Thus, the elevated vesicle number likely corresponds to the increased uptake of ACs, rather than the blockage in phagosome maturation (Kurant et al. 2008; Petrignani et al. 2021).

These findings highlight a functional distinction between NimB1 and NimB4, with NimB1 regulating the early stages of apoptotic cell uptake and NimB4 playing a critical role in the later steps of lysosomal fusion and degradation (Petrignani et al. 2021).

## Discussion

Bridging molecules play a critical role in apoptotic cell clearance by linking phagocytes to their targets (Hanayama et al. 2002; Elliott and Ravichandran 2016; Petrignani et al. 2021; Ji et al. 2023). Previous studies in *Drosophila* have identified NimB4 and Orion as essential mediators of efferocytosis. While NimB4 facilitates phagosome maturation and lysosomal fusion, Orion is involved in the early recognition phase of phagocytosis, facilitating the detection of “eat-me” signals by enhancing PS binding to the transmembrane receptor Draper (Ji et al. 2023; Petrignani et al. 2021). These roles parallel mammalian systems, where proteins like MFG-E8 link PS to integrins to facilitate apoptotic cell engulfment (Hanayama et al. 2002). Our study identifies NimB1 as a new player in efferocytosis, with a distinct role in down-regulating early phagocytic uptake, and illustrating how apoptotic cell clearance is fine-tuned at multiple stages of the phagocytic process.

Our study reveals striking similarities between NimB1 and NimB4. Phylogenetic and structural analyses support the hypothesis that NimB1 and NimB4 emerged from a recent duplication event in the NimB family, with NimB2 representing an earlier evolutionary divergence. NimB1 and NimB4 share conserved PS-binding motifs and exhibit significant structural overlap, including the presence of PS-binding pockets. Functional assays demonstrate that similar to NimB4, NimB1 binds apoptotic corpses in a PS-dependent manner. However, in contrast to the delayed maturation and phagolysosome accumulation observed in *NimB4* mutants, *NimB1* mutants exhibit accelerated apoptotic cell uptake. These findings suggest that NimB1 and NimB4 have opposing roles, where NimB1 delays early phagocytic events while NimB4 promotes downstream lysosome fusion. Phagocytosis of ACs must be tightly regulated as accumulation of apoptotic debris due to either a delay in processing or accelerated uptake may impair hemocyte function (Freeman et al. 2003; Kinchen et al. 2008; Kurant et al. 2008; Petrignani et al. 2021). We speculate that NimB1 and NimB4 have evolved to fine-tune the rate of efferocytosis, notably at the embryonic stage when the tissue renewal rate is high, and these proteins peak in expression.

This division of labor in efferocytosis is mirrored in the complementary functions of Draper isoforms (Logan et al. 2012). Draper, a homolog of the mammalian MEGF10 and CED-1, has three isoforms with distinct roles in phagocytosis. Draper I features an immunoreceptor tyrosine-based activation motif (ITAM) that promotes recognition and internalization of ACs by initiating signaling cascades. In contrast, Draper II contains an immunoreceptor tyrosine-based inhibition motif (ITIM), which down-modulates the phagocytic response post-engulfment and regulates the progression of phagocytosis. The function of Draper III is not fully understood. This balance between activation by Draper I and inhibition by Draper II is thought to prevent overactivation of phagocytes, which could lead to phagocyte impairment or immune dysregulation during periods of high apoptotic turnover, such as development or tissue repair (Logan et al. 2012). The involvement of Draper I in engulfment mirrors the role of NimB4 in facilitating maturation and lysosomal fusion, while the inhibitory functions of Draper II parallel modulation of early recognition and binding events by NimB1. Further study should investigate whether the effects of NimB1 and NimB4 are mediated via Draper isoforms. The cooperative roles of the Draper isoforms and NimB proteins illustrate a complex regulatory network that optimizes apoptotic cell clearance by *Drosophila* phagocytes.

The combinatory phenotype of *NimB1, NimB4* double mutants further underscores the complementary roles of these genes, with double mutants displaying both the enhanced cell surface remodeling observed in the *NimB1* mutant and accumulation of unresolved phagosomes observed in *NimB4* or *draper* null mutants. Overall, phagocytic defect observed in double mutants is very similar to the *NimB4* single mutant. This suggests that the function of NimB4 in phagosome maturation is dominant, echoing the observation that the *draper* mutant lacking both Draper I and II isoforms has a phenotype similar to a mutant of Draper I alone (Logan et al. 2012). This consistent with a scenario in which NimB1 and Draper II antagonize the effects of NimB4 and Draper I. AlphaFold2 modeling predicts a strong interaction interface between NimB1 and NimB4, raising the possibility of direct contact between the two factors. Experimental validation of this interaction would provide valuable insights into the molecular mechanisms underlying their complementary functions.

Finally, the evolutionary specialization of the Nimrod family exemplifies how gene duplication and divergence can create adaptive strategies for fine tuning immune regulation. By partitioning tasks between proteins that delay early recognition (NimB1) and promote late-stage processing (NimB4), a balance is achieved that ensures efficient apoptotic clearance without compromising phagocyte viability. Similar strategies are observed in mammals, where proteins like CD47 and TIM-4 maintain immune homeostasis through opposing functions in phagocytosis (Elliott and Ravichandran 2016). *NimB1* mutants appear to be viable, while *NimB4* like *draper* mutants display reduced viability and neurodegeneration. It is likely that the role of NimB1 is subtler but may become apparent in more critical situations such as recurrent waves of apoptosis.

In conclusion, our study identifies NimB1 as a novel modulator of early apoptotic cell recognition and internalization in *Drosophila*, complementing and opposing the role of NimB4 in facilitating phagosome maturation. In addition to this study of NimB1, recent studies of Orion and NimB4 reveal the role of secreted factors in *Drosophila* efferocytosis, similar to the bridging molecules of mammals. Future studies should explore the interactions between the secreted factors NimB1, NimB4, Orion and Draper isoforms as well as SIMU receptor, and further explore the conserved principles of apoptotic cell clearance across species.

## Materials and methods

### *Drosophila* stocks and rearing conditions

All *Drosophila* stocks were maintained at 25°C on standard fly medium consisting of 6% cornmeal, 6% yeast, 0.6% agar, 0.1% fruit juice (consisting of 50% grape juice and 50% multifruit/ multivitamin juice), supplemented with 10.6 g.L^-1^ moldex and 4.9 mL.L^-1^ propionic acid. Third instar (L3) wandering larvae were selected at 110-120 h AEL. The following fly lines were used in this study:

### Mutant and transgenic line generation

*NimB1^229^* flies were generated as follows: *w^iso^*embryos were injected with a mixture of recombinant *Cas9* (Invitrogen) and a gRNA targeting the *NimB1* coding sequence. The following gRNA sequence was used: *TATGTCGCCGTAGAGCTCCGTGG*. The mutant candidates derived from male parents carrying *nos-Cas9* and *U6-gRNA* transgenes were screened by PCR and direct sequencing of the target region. gRNA was synthesized *in vitro* using the GeneArt precision synthesis kit (ThermoFisher) and purified following the manufacturer’s instructions. 300ng of *NimB1* gRNA and 300ng of *ebony* gRNA were mixed with 4.5 μg of GeneArt Cas9 Platinum (ThermoFisher) in 10mM Tris pH7.8, incubated 15min at 37°C and microinjected into *w^1118^* embryos. G_0_ ebony^-^ mosaics flies were selected and crossed to *w^1118^;TM6c/Xa* flies and F_1_ were screened by T7 endonuclease assay (B. C. Lee et al. 2014; Rommelaere et al. 2019). *NimB1^229^* mutant into the *w^1118^*DrosDel isogenic background for seven generations.

To generate the *NimB1^61^,NimB4^sk2^* double mutant, CRISPR-Cas9 was also employed to target the *NimB1* locus in *NimB4^sk2^* isogenic mutant background. A 14-bp deletion was introduced in exon 1 of *NimB1*, leading to a frameshift mutation and premature stop codon. The double mutant combined the *NimB4^sk2^* mutation with the *NimB1^61^* mutation. The double mutant was verified by PCR genotyping and Sanger sequencing. This double mutant was backcrossed into the *w^1118^* DrosDel isogenic background for at least seven generations to ensure a consistent genetic background for all experiments.

For the *UAS-NimB1-RFP* line, the *NimB1* cDNA sequence without STOP codon was cloned into the entry vector pENTR/D-Topo (Invitrogen) and subsequently shuttled into the RFP expression vectors pTWR (C-terminal RFP tag), obtained from the DGRC *Drosophila* Gateway vector collection. Plasmids were injected in house.

### Tools validation

Using qPCR, we validated the transcriptional activation of *UAS-NimB1* and *UAS-NimB1-RFP* with both *Hml^p2a^-Gal4* and ubiquitous *Actin^5c^-Gal4* drivers. These drivers effectively induced NimB1 overexpression, but RNAi-mediated knockdown using *NimB1-RNAi-TRIP* (Bloomington) and *NimB1-RNAi-KK* (Vienna) failed to achieve significant silencing.

To confirm NimB1 expression at the protein level, we performed Western blot analysis on hemolymph extracts from third instar larvae. We detected NimB1-RFP at its expected molecular weight of 44 kDa, verifying its secretion into the hemolymph (Supplementary Document S3 C).

### RT–qPCR experiments

For quantification of mRNA, whole third instar larvae (n=15) or dissected tissues (n=20-40) were isolated by TRIzol reagent and dissolved in RNase-free water. 500ng total RNA was then reverse-transcribed in 10µL reaction volume using PrimeScript RT (TAKARA) and a mixture of oligo-dT and random hexamer primers. Quantitative PCR was performed on cDNA samples on a LightCycler 480 (Roche) in 96-well plates using the Quant Studio™ (Applied Biosystems™) with SYBR™ Select Master (Applied Biosystems™). Expression values were normalized to *RpL32* or *Rp49*. Primer sequences used are provided in Table S2.

### Western blot

Hemolymph samples were collected as follows: forty third instar larvae were bled on a glass slide on ice, mixed with 10 μL of PBS supplemented with complete protease inhibitor solution (Roche) and 1mM phenylmethylsulfonyl fluoride (Sigma) and N-Phenylthiourea (Sigma), and then centrifuged for 10 minutes at 1’000 g, 4°C. This was followed by a second centrifugation for 5 min at 15 000 g. Protein concentration of the samples was determined by BCA assay and 40 μg of protein extract was separated on a 4%–12% acrylamide precast Novex NuPage gel (Invitrogen) under reducing conditions and transferred to membranes (Invitrogen iBlot 2). After blocking in 5% non-fat dry milk in PBS containing 0.1% Tween for 1h, membranes were incubated at 4°C overnight with a mouse anti-RFP antibody (Abcam) in a 1:1000 dilution. Anti-mouse-HRP secondary antibody (Jackson ImmunoResearch) in a 1:15’000 dilution was incubated for 45 min at room temperature. Bound antibody was detected using ECL (GE Healthcare) according to the manufacturer’s instructions. Membranes were imaged on a ChemiDoc XRS+ (Biorad).

### Apoptotic cell preparation

The S2 cells were cultured in Schneider’s insect medium (Sigma-Aldrich) containing 10% FBS (Gibco™), penicillin (Sigma-Aldrich) and streptomycin (Sigma-Aldrich) at a concentration of 100 U/ml. To induce apoptosis, cycloheximide (CHX, Sigma-Aldrich) was added at a final concentration of 50μg/ml. 24 hours after the cycloheximide treatment, the cells were isolated and removed by pelleting with centrifugation at 400g for 5 min at 4°C. In order to stain the apoptotic bodies, CellTrace™ CFSE Cell Proliferation Kit (Invitrogen^TM^), for flow cytometry (or CellTracker^TM^ Red CMTPX Dye) was added to the supernatant harvested from the cells at a final concentration of 5 μM and incubated 15 min at room temperature in the dark.

### Immunohistochemistry

For immunofluorescence, third instar larvae were dissected into 150 μL PBS pH 7.4, and macrophages were allowed to adhere on a glass slide for 40 min and fixed for 10 min in PBS containing 4% paraformaldehyde. For immunostaining, fixed cells were subsequently rinsed in PBS + 0.1% Triton X-100 (PBT), permeabilized and blocked in PBT + 2% bovine serum albumin (BSA) for 1h and incubated with primary antibodies in PBT + 2% BSA overnight at 4°C. After 1h washing, secondary antibodies and DAPI were applied at room temperature for 2h. Primary antibodies are as follows: chicken anti-GFP (Abcam, 1:1,000), rabbit anti-SIMU 1:1000 (Shklyar et al. 2013; Shklover, Levy-Adam, and Kurant 2015; Kurant et al. 2008), rabbit anti RFP (cell Signaling, 1:1000), Rab7 (AB 2722471 DSHB, 1:500). Alexa488 and Alexa555-conjugated secondary antibodies (Life technologies, 1:2000) were used.

### Macrophages LysoTracker red staining

Macrophages were allowed to adhere to slides for 45 min, then incubated with 1 μM LysoTracker® Red DND-99 (Invitrogen^TM^, L7528) in PBS for 1 min at RT. The samples were washed twice in PBS and mounted for immediate observations under epifluorescence or confocal microscope.

### Transmission electron microscopy

Third instar wandering larvae were bled in 50μL of Schneider’s insect medium (Sigma-Aldrich). The collected hemolymph was incubated on a glass coverslip for 1h before being fixed for 2h with 2% paraformaldehyde + 2.5% glutaraldehyde in 0.1 M phosphate buffer, pH 7.4. Samples were then washed in cacodylate buffer (0.1 M, pH 7.4), fixed again in 1% osmium tetroxide and potassium ferrocyanide 1.5% in cacodylate buffer. After several washes in distilled water, samples were stained in 1% uranyl acetate in water, washed again, and then dehydrated in graded alcohol series (50%, 70%, 90%, 95%, 100%). Embedding was performed first in 1:1 Hard EPON and ethanol 100%, and afterwards in pure EPON, before being embedded on coated glass slides and placed at 60 °C overnight. Images were acquired with a FEI Tecnai Spirit 120 kV (FEI Company, Eagle, The Netherlands).

### Protein modeling

AlphaFold2 was employed to model the structures of NimB1, NimB2 and NimB4. The result structures were visualized and analyzed using PyMOL software for alignment and superposition for identifying the conserved domains. The RRY motifs, identified as conserved PS-binding regions based on alignment with the Orion study (Hu et al., 2022), were visualized using PyMOL (company). The indices used to assess the similarity between the proteins were quantified using the TM align method (Zhang and Skolnick 2005) by alignment length, Root-Mean Square Deviation (RMSD), Sequence identity, and the TM-score.

### Binding assay with ACs

Third instar wandering larvae were bled into Schneider’s insect medium (Sigma-Aldrich) on a previously chilled glass slide. After larva bleeding, macrophages and ACs (CellTracker^TM^ Red CMTPX Dye) were incubated directly on the prechilled slide, in cold Schneider’s medium, on ice for 60 min. After fixation in 4% paraformaldehyde PBS, AlexaFluor488-phalloidin staining (Molecular Probes^TM^) was performed. Finally, cells were stained with 1:15 000 dilution of DAPI (Sigma-Aldrich). Fluorescent preparations were mounted in Dako fluorescence media.

### *Ex vivo* larval macrophage phagocytosis assay

Five third instar wandering larvae were bled into 150 μL of Schneider’s insect medium (Sigma-Aldrich). The macrophage suspension was then transferred to 1.5 mL low binding tubes (LoBind, Eppendorf, Hamburg, Germany). The samples were incubated respectively with 2×10^7^ Bioparticles^®^ conjugate from *Staphylococcus aureus* (Wood strain without protein A) BioParticles™, Alexa Fluor™ 488 conjugate (Invitrogen™) or *Escherichia coli* (K-12 strain) BioParticles™, Alexa Fluor™ 488 (Invitrogen™) conjugate for 60 or 120 min to enable phagocytosis and then placed on ice to stop the reaction. CellTrace™ CFSE Cell Proliferation Kit (Invitrogen™) was used to stain S2 ACs prepared as mentioned above and incubated with dissected macrophages for 60 or 120 min for flow cytometry. The concentration of ACs was normalized to the amount of macrophages in the probe counting to 10-15 ACs per macrophage. The stained ACs were measured after preparation and diluted to the desired concentration prior to the experiment. Phagocytosis was quantified using a flow cytometer (Cytoflex, Beckman Counter) to measure the fraction of cells phagocytosing, and their fluorescence intensity. 60 μL volume was read at medium speed (30 μL/min). In a first step, macrophages were identified using the *HmlΔ-Gal4,UAS-GFP* live staining. The fluorescence intensity of single macrophages was measured in the green channel with 488nm laser and 530/30 standard filter. At least 2000 cells per genotype and per assay were analyzed. The results are an average of three independent experiments.

The phagocytic index was calculated as follows:

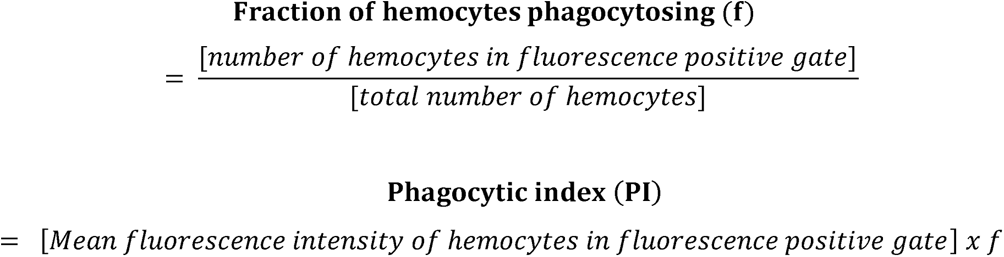

For the short time point phagocytosis the procedure was followed as for the above.

### *In vivo* pinching assay

Mechanical induction of apoptosis was performed on third instar larvae by pinching the cuticle near the posterior end with forceps. Larvae were maintained at 25°C on food for 90 min post-pinching to allow apoptotic cell generation. Macrophages were isolated and stained with DAPI. Samples were imaged using confocal microscopy to assess apoptosis induction. The apoptotic events were quantified as additional DAPI events with DAPI nucleus stainings excluded. A Leica Sp8 microscope was used for image acquisition. The quantification method used Fiji software for Z stack projections, channel separation, and brightness and contrast adjustments. QuPath and Cell pose software were used to subtract the nucleus and to expose only the cytosolic events for quantification.

### Image analysis and quantification

All images used for quantification were captured with a Leica SP8 microscope, and all analyses were performed using ImageJ. For quantification of the fluorescence signal intensity, the fluorescent images were first converted to 8-bit images, and the total intensity value with an identical threshold was captured and measured with ImageJ. The freehand selection tool in ImageJ was used to capture and measure the area of the macrophages. Colocalization analysis was done with the ImageJ plugin ‘Just another Co-localization Plugin’ after channel splitting and background subtraction. Rr (Pearson’s correlation coefficient), Ch1:Ch2 ratios, M1 and M2 (Manders’ colocalization coefficient for channel 1 and 2) were tabulated for each image.

### Statistical tests

Each experiment was repeated independently a minimum of three times (unless otherwise indicated), error bars represent the standard deviation (SD) of replicate experiments. Data were analyzed using appropriate statistical tests as indicated in figure legends using the GraphPad Prism software. Significance tests were performed using the Mann-Whitney test. For experiments with more than 2 conditions, significance was tested using ANOVA test (with a confidence level of 95%). P values of < 0.05 = *, < 0.01 = **, and < 0.001 = ***.

## Acknowledgments

We thank the VDRC in Vienna, the Bloomington Stock Center, Luis Teixeira, Marc Freeman, Jose C. Pastor-Pareja and Holy Stephenson for the fly stocks. We thank the BIOP platform (EPFL) and the BioEM platform (EPFL) for help with electron microscopy experiments. We thank Hannah Westlake for critical reading of the manuscript and Lukas Neukomm and Brian McCabe for stimulating discussions.

## Author contributions

AD, SR, EK and BL conceived and designed the experiments and wrote the paper. AD and AG performed the experiments. AD, SR, EK analyzed the data. AD and BL wrote the manuscript.

---

**Table S1.**
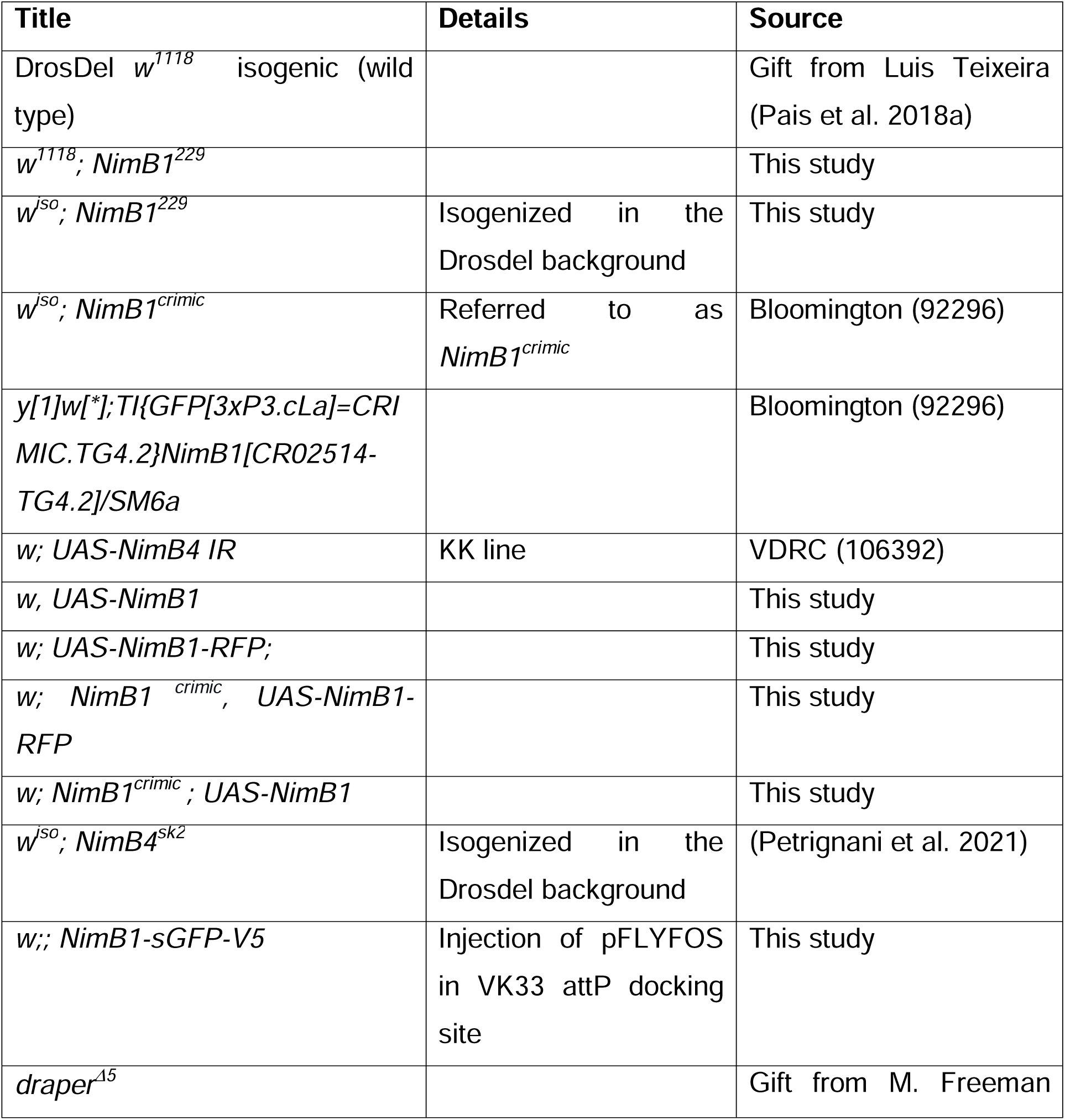
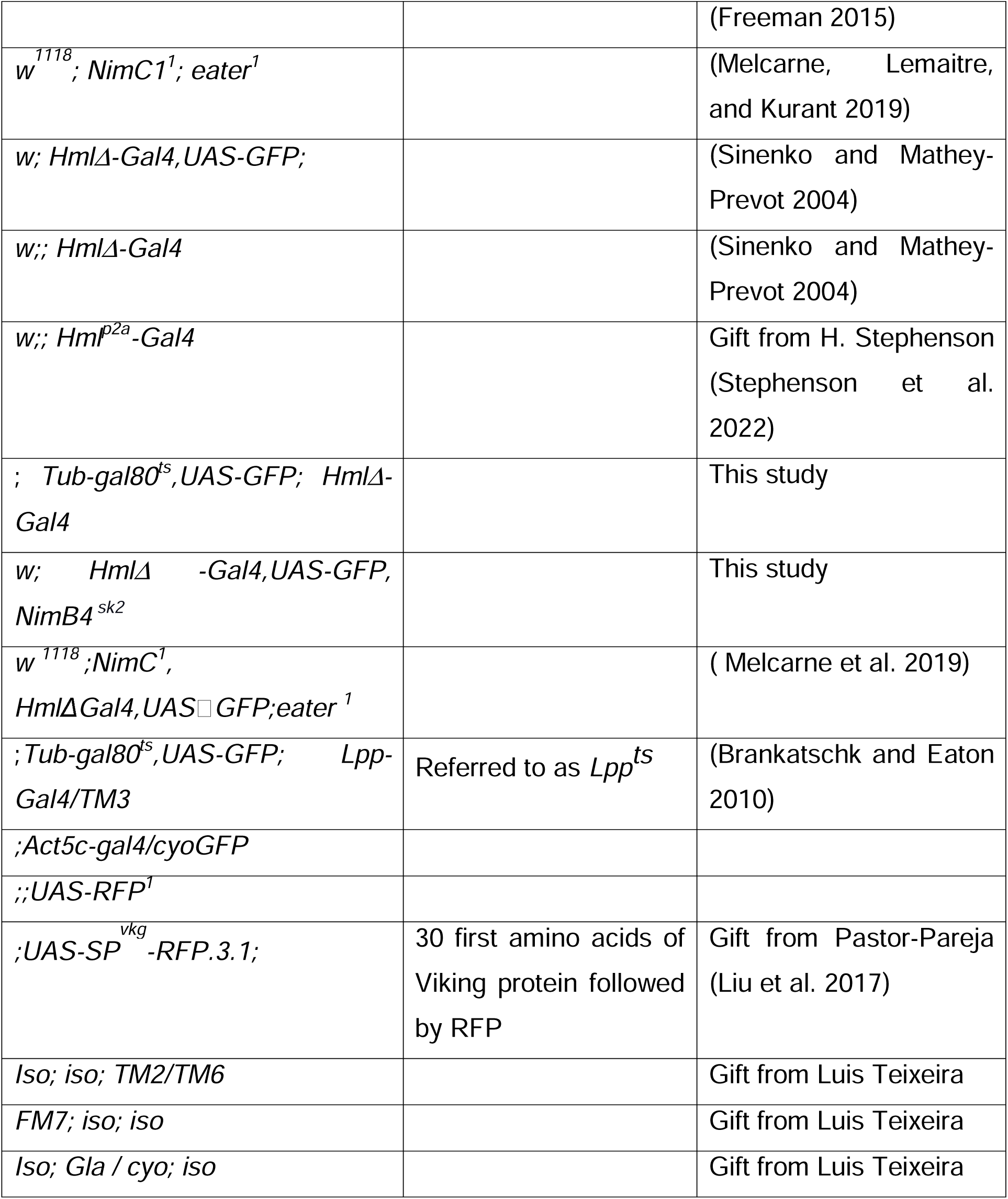

**Table S2.**
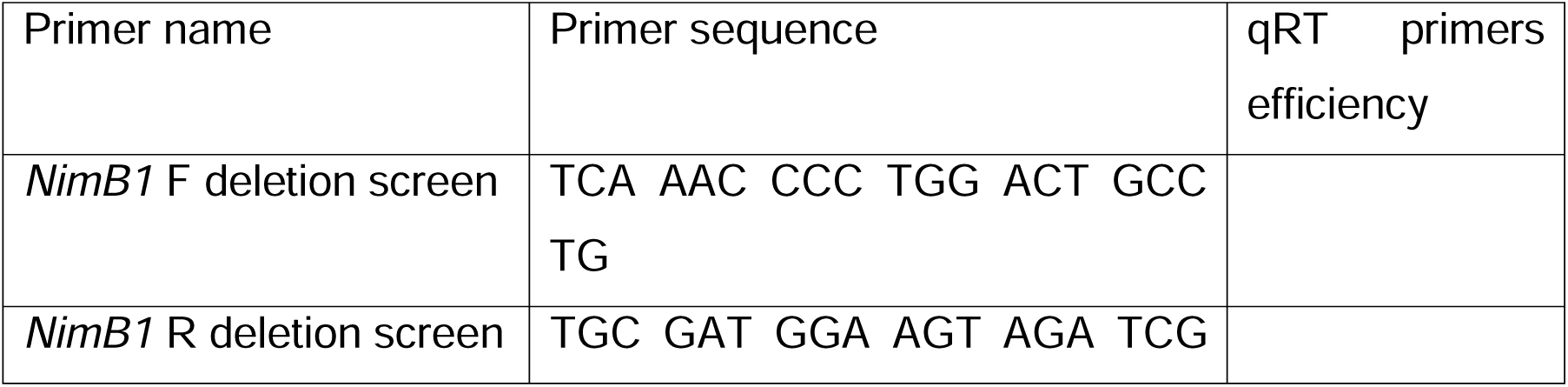
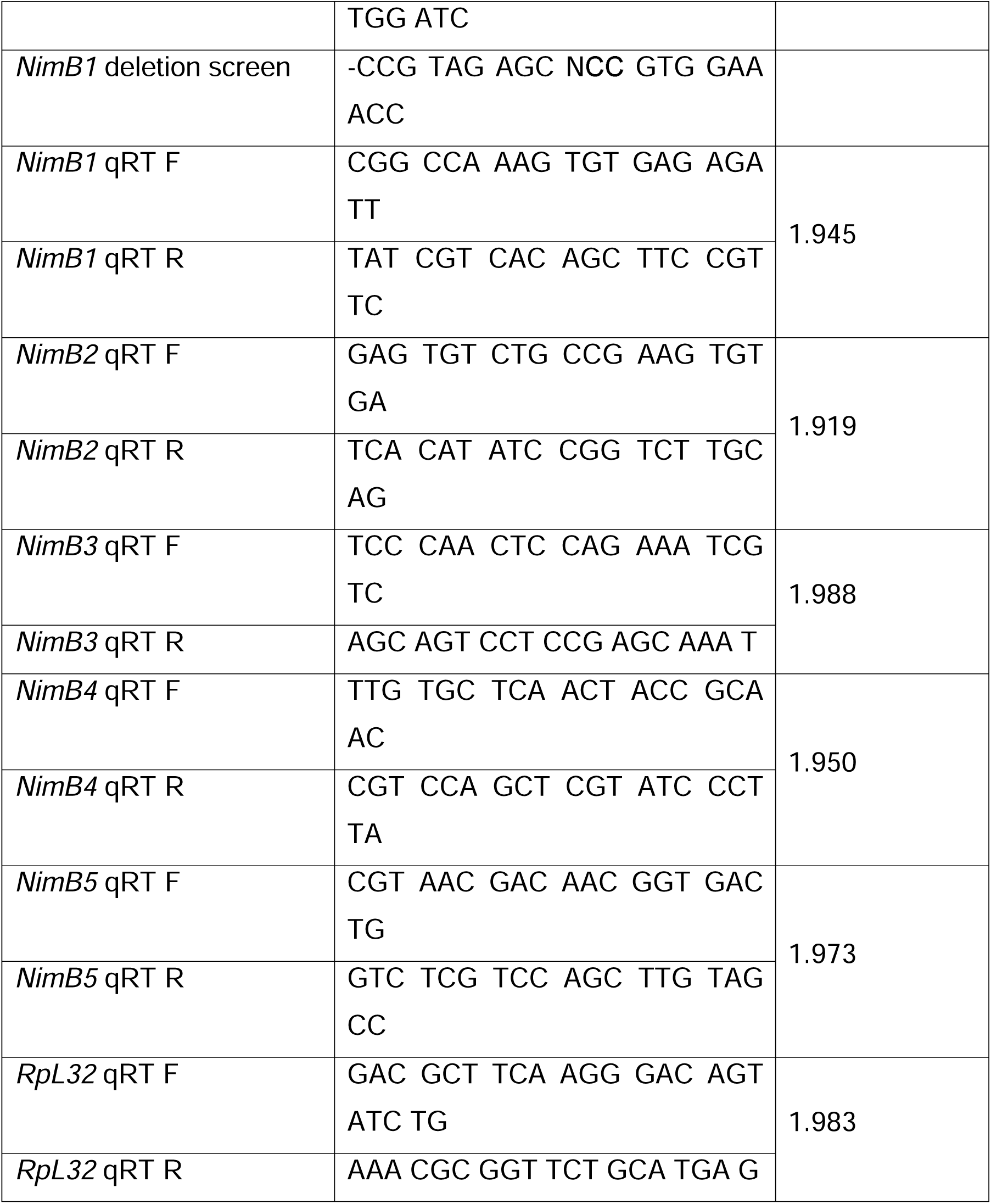

## Bibliography

Akakura, Shin, Sukhwinder Singh, Matthew Spataro, Reiko Akakura, Jong-Il Kim, Matthew L. Albert, and Raymond B. Birge. 2004. ‘The Opsonin MFG-E8 Is a Ligand for the Alphavbeta5 Integrin and Triggers DOCK180-Dependent Rac1 Activation for the Phagocytosis of Apoptotic Cells’. Experimental Cell Research 292 (2): 403–16. 10.1016/j.yexcr.2003.09.011.

Arandjelovic, Sanja, and Kodi S. Ravichandran. 2015. ‘Phagocytosis of Apoptotic Cells in Homeostasis’. Nature Immunology 16 (9): 907–17. 10.1038/ni.3253.

Barth, Nicole D., John A. Marwick, Marc Vendrell, Adriano G. Rossi, and Ian Dransfield. 2017. ‘The “Phagocytic Synapse” and Clearance of Apoptotic Cells’. Frontiers in Immunology 8 (December):1708. 10.3389/fimmu.2017.01708.

Boada-Romero, Emilio, Jennifer Martinez, Bradlee L. Heckmann, and Douglas R. Green. 2020. ‘The Clearance of Dead Cells by Efferocytosis’. Nature Reviews. Molecular Cell Biology 21 (7): 398–414. 10.1038/s41580-020-0232-1.

Borisenko, G. G., S. L. Iverson, S. Ahlberg, V. E. Kagan, and B. Fadeel. 2004. ‘Milk Fat Globule Epidermal Growth Factor 8 (MFG-E8) Binds to Oxidized Phosphatidylserine: Implications for Macrophage Clearance of Apoptotic Cells’. Cell Death & Differentiation 11 (8): 943–45. 10.1038/sj.cdd.4401421.

Bork, P., A. K. Downing, B. Kieffer, and I. D. Campbell. 1996. ‘Structure and Distribution of Modules in Extracellular Proteins’. Quarterly Reviews of Biophysics 29 (2): 119–67. 10.1017/s0033583500005783.

Brankatschk, Marko, and Suzanne Eaton. 2010. ‘Lipoprotein Particles Cross the Blood-Brain Barrier in Drosophila’. The Journal of Neuroscience: The Official Journal of the Society for Neuroscience 30 (31): 10441–47. 10.1523/JNEUROSCI.5943-09.2010.

Bretscher, Andrew J., Viktor Honti, Olivier Binggeli, Olivier Burri, Mickael Poidevin, Éva Kurucz, János Zsámboki, István Andó, and Bruno Lemaitre. 2015. ‘The Nimrod Transmembrane Receptor Eater Is Required for Hemocyte Attachment to the Sessile Compartment in Drosophila Melanogaster’. Biology Open 4 (3): 355–63. 10.1242/bio.201410595.

Brown, Guy C. 2024. ‘Cell Death by Phagocytosis’. Nature Reviews Immunology 24 (2): 91–102. 10.1038/s41577-023-00921-6.

Callahan, M. K., P. Williamson, and R. A. Schlegel. 2000. ‘Surface Expression of Phosphatidylserine on Macrophages Is Required for Phagocytosis of Apoptotic Thymocytes’. Cell Death & Differentiation 7 (7): 645–53. 10.1038/sj.cdd.4400690.

Cockram, Tom O. J., Jacob M. Dundee, Alma S. Popescu, and Guy C. Brown. 2021. ‘The Phagocytic Code Regulating Phagocytosis of Mammalian Cells’. Frontiers in Immunology 12:629979. 10.3389/fimmu.2021.629979.

Davidson, Andrew J., and Will Wood. 2020. ‘Macrophages Use Distinct Actin Regulators to Switch Engulfment Strategies and Ensure Phagocytic Plasticity In Vivo’. Cell Reports 31 (8): 107692. 10.1016/j.celrep.2020.107692.

Elliott, Michael R., and Kodi S. Ravichandran. 2008. ‘Death in the CNS: Six-Microns-Under’. Cell 133 (3): 393–95. 10.1016/j.cell.2008.04.014.

Elliott, Michael R., and Kodi S. Ravichandran. 2016. ‘The Dynamics of Apoptotic Cell Clearance’. Developmental Cell 38 (2): 147–60. 10.1016/j.devcel.2016.06.029.

Engeland, M. van, L. J. Nieland, F. C. Ramaekers, B. Schutte, and C. P. Reutelingsperger. 1998. ‘Annexin V-Affinity Assay: A Review on an Apoptosis Detection System Based on Phosphatidylserine Exposure’. Cytometry 31 (1): 1–9. 10.1002/(sici)1097-0320(19980101)31:1<1::aid-cyto1>3.0.co;2-r.

Fadok, V. A., D. L. Bratton, A. Konowal, P. W. Freed, J. Y. Westcott, and P. M. Henson. 1998. ‘Macrophages That Have Ingested Apoptotic Cells in Vitro Inhibit Proinflammatory Cytokine Production through Autocrine/Paracrine Mechanisms Involving TGF-Beta, PGE2, and PAF’. The Journal of Clinical Investigation 101 (4): 890–98. 10.1172/JCI1112.

Flannagan, Ronald S., Valentin Jaumouillé, and Sergio Grinstein. 2012. ‘The Cell Biology of Phagocytosis’. Annual Review of Pathology 7:61–98. 10.1146/annurev-pathol-011811-132445.

Freeman, Marc R. 2015. ‘Drosophila Central Nervous System Glia’. Cold Spring Harbor Perspectives in Biology 7 (11): a020552. 10.1101/cshperspect.a020552.

Freeman, Marc R., Jeffrey Delrow, Junhyong Kim, Eric Johnson, and Chris Q. Doe. 2003. ‘Unwrapping Glial Biology: Gcm Target Genes Regulating Glial Development, Diversification, and Function’. Neuron 38 (4): 567–80. 10.1016/s0896-6273(03)00289-7.

Fullard, John F., Abhijit Kale, and Nicholas E. Baker. 2009. ‘Clearance of Apoptotic Corpses’. Apoptosis: An International Journal on Programmed Cell Death 14 (8): 1029–37. 10.1007/s10495-009-0335-9.

Fuller, Abby D., and Linda J. Van Eldik. 2008. ‘MFG-E8 Regulates Microglial Phagocytosis of Apoptotic Neurons’. Journal of Neuroimmune Pharmacology: The Official Journal of the Society on NeuroImmune Pharmacology 3 (4): 246–56. 10.1007/s11481-008-9118-2.

Hamon, Yannick, Doriane Trompier, Zhong Ma, Victor Venegas, Matthieu Pophillat, Vincent Mignotte, Zheng Zhou, and Giovanna Chimini. 2006. ‘Cooperation between Engulfment Receptors: The Case of ABCA1 and MEGF10’. PloS One (1): e120. 10.1371/journal.pone.0000120.

Hanayama, Rikinari, Masato Tanaka, Keiko Miwa, Azusa Shinohara, Akihiro Iwamatsu, and Shigekazu Nagata. 2002. ‘Identification of a Factor That Links Apoptotic Cells to Phagocytes’. Nature 417 (6885): 182–87. 10.1038/417182a.

Hashimoto, Yumi, Yukichika Tabuchi, Kenji Sakurai, Mayumi Kutsuna, Kenji Kurokawa, Takeshi Awasaki, Kazuhisa Sekimizu, Yoshinobu Nakanishi, and Akiko Shiratsuchi. 2009. ‘Identification of Lipoteichoic Acid as a Ligand for Draper in the Phagocytosis of Staphylococcus Aureus by Drosophila Hemocytes’. Journal of Immunology (Baltimore, Md.: 1950) 183 (11): 7451–60. 10.4049/jimmunol.0901032.

Hilu-Dadia, Reut, Aseel Ghanem, Shelly Vogelesang, Malak Ayoub, Ketty Hakim-Mishnaevski, and Estee Kurant. 2025. ‘Santa-Maria Is a Glial Phagocytic Receptor That Acts with SIMU to Recognize and Engulf Apoptotic Neurons’. Cell Report in Press.

Ivy, Jessica R., Maik Drechsler, James H. Catterson, Rolf Bodmer, Karen Ocorr, Achim Paululat, and Paul S. Hartley. 2015. ‘Klf15 Is Critical for the Development and Differentiation of Drosophila Nephrocytes’. PLOS ONE 10 (8): e0134620. 10.1371/journal.pone.0134620.

Janko, Christina, Ivica Jeremic, Mona Biermann, Ricardo Chaurio, Christine Schorn, Luis E. Muñoz, and Martin Herrmann. 2013. ‘Cooperative Binding of Annexin A5 to Phosphatidylserine on Apoptotic Cell Membranes’. Physical Biology 10 (6): 065006. 10.1088/1478-3975/10/6/065006.

Ji, Hui, Bei Wang, Yifan Shen, David Labib, Joyce Lei, Xinchen Chen, Maria Sapar, Ana Boulanger, Jean-Maurice Dura, and Chun Han. 2023. ‘The Drosophila Chemokine–like Orion Bridges Phosphatidylserine and Draper in Phagocytosis of Neurons’. Proceedings of the National Academy of Sciences 120 (24): e2303392120. 10.1073/pnas.2303392120.

Ke, Hongmei, Zhi Feng, Min Liu, Tianhui Sun, Jianli Dai, Mengqi Ma, Lu-Ping Liu, Jian-Quan Ni, and José Carlos Pastor-Pareja. 2018. ‘Collagen Secretion Screening in *Drosophila* Supports a Common Secretory Machinery and Multiple Rab Requirements’. Journal of Genetics and Genomics 45 (6): 299–313. 10.1016/j.jgg.2018.05.002.

Kinchen, Jason M., Kimon Doukoumetzidis, Johann Almendinger, Lilli Stergiou, Annie Tosello-Trampont, Costi D. Sifri, Michael O. Hengartner, and Kodi S. Ravichandran. 2008. ‘A Pathway for Phagosome Maturation during Engulfment of Apoptotic Cells’. Nature Cell Biology 10 (5): 556–66. 10.1038/ncb1718.

Kocks, Christine, Ju Hyun Cho, Nadine Nehme, Johanna Ulvila, Alan M. Pearson, Marie Meister, Charles Strom, et al. 2005. ‘Eater, a Transmembrane Protein Mediating Phagocytosis of Bacterial Pathogens in Drosophila’. Cell 123 (2): 335–46. 10.1016/j.cell.2005.08.034.

Koopman, G., C. P. Reutelingsperger, G. A. Kuijten, R. M. Keehnen, S. T. Pals, and M. H. van Oers. 1994. ‘Annexin V for Flow Cytometric Detection of Phosphatidylserine Expression on B Cells Undergoing Apoptosis’. Blood 84 (5): 1415–20.

Kurant, Estee, Sofia Axelrod, Dan Leaman, and Ulrike Gaul. 2008. ‘Six-Microns-Under Acts Upstream of Draper in the Glial Phagocytosis of Apoptotic Neurons’. Cell 133 (3): 498–509. 10.1016/j.cell.2008.02.052.

Kurucz, Eva, Róbert Márkus, János Zsámboki, Katalin Folkl-Medzihradszky, Zsuzsanna Darula, Péter Vilmos, Andor Udvardy, et al. 2007. ‘Nimrod, a Putative Phagocytosis Receptor with EGF Repeats in Drosophila Plasmatocytes’. Current Biology: CB 17 (7): 649–54. 10.1016/j.cub.2007.02.041.

Kusunoki, Ryusaku, Shunji Ishihara, Monowar Aziz, Akihiko Oka, Yasumasa Tada, and Yoshikazu Kinoshita. 2012. ‘Roles of Milk Fat Globule-Epidermal Growth Factor 8 in Intestinal Inflammation’. Digestion 85 (2): 103–7. 10.1159/000334679.

Lee, Byung Cheon, Alaattin Kaya, Siming Ma, Gwansu Kim, Maxim V. Gerashchenko, Sun Hee Yim, Zhen Hu, Lawrence G. Harshman, and Vadim N. Gladyshev. 2014. ‘Methionine Restriction Extends Lifespan of Drosophila Melanogaster under Conditions of Low Amino-Acid Status’. Nature Communications 5 (April):3592. 10.1038/ncomms4592.

Lee, Pei-Tseng, Jonathan Zirin, Oguz Kanca, Wen-Wen Lin, Karen L. Schulze, David Li-Kroeger, Rong Tao, et al. 2018. ‘A Gene-Specific T2A-GAL4 Library for Drosophila’. eLife 7 (March):e35574. 10.7554/eLife.35574.

Lemke, Greg. 2019. ‘How Macrophages Deal with Death’. Nature Reviews Immunology 19 (9): 539–49. 10.1038/s41577-019-0167-y.

Liu, Min, Zhi Feng, Hongmei Ke, Ying Liu, Tianhui Sun, Jianli Dai, Wenhong Cui, and José Carlos Pastor-Pareja. 2017. ‘Tango1 Spatially Organizes ER Exit Sites to Control ER Export’. The Journal of Cell Biology 216 (4): 1035–49. 10.1083/jcb.201611088.

Logan, Mary A., Rachel Hackett, Johnna Doherty, Amy Sheehan, Sean D. Speese, and Marc R. Freeman. 2012. ‘Negative Regulation of Glial Engulfment Activity by Draper Terminates Glial Responses to Axon Injury’. Nature Neuroscience 15 (5): 722–30. 10.1038/nn.3066.

MacDonald, Jennifer M., Margaret G. Beach, Ermelinda Porpiglia, Amy E. Sheehan, Ryan J. Watts, and Marc R. Freeman. 2006. ‘The Drosophila Cell Corpse Engulfment Receptor Draper Mediates Glial Clearance of Severed Axons’. Neuron 50 (6): 869–81. 10.1016/j.neuron.2006.04.028.

Manaka, Junko, Takayuki Kuraishi, Akiko Shiratsuchi, Yuji Nakai, Haruhiro Higashida, Peter Henson, and Yoshinobu Nakanishi. 2004. ‘Draper-Mediated and Phosphatidylserine-Independent Phagocytosis of Apoptotic Cells by Drosophila Hemocytes/Macrophages’. The Journal of Biological Chemistry 279 (46): 48466–76. 10.1074/jbc.M408597200.

Mangahas, Paolo M., and Zheng Zhou. 2005. ‘Clearance of Apoptotic Cells in Caenorhabditis Elegans’. Seminars in Cell & Developmental Biology 16(2): 295–306. 10.1016/j.semcdb.2004.12.005.

Melcarne, C., B. Lemaitre, and E. Kurant. 2019. ‘Phagocytosis in Drosophila: From Molecules and Cellular Machinery to Physiology’. Insect Biochemistry and Molecular Biology 109 (June):1–12. 10.1016/j.ibmb.2019.04.002.

Melcarne, Claudia, Elodie Ramond, Jan Dudzic, Andrew J. Bretscher, Éva Kurucz, István Andó, and Bruno Lemaitre. 2019. ‘Two Nimrod Receptors, NimC1 and Eater, Synergistically Contribute to Bacterial Phagocytosis in Drosophila Melanogaster’. The FEBS Journal 286 (14): 2670–91. 10.1111/febs.14857.

Nagaosa, Kaz, Ryo Okada, Saori Nonaka, Kazuki Takeuchi, Yu Fujita, Tomoyuki Miyasaka, Junko Manaka, István Ando, and Yoshinobu Nakanishi. 2011. ‘Integrin Βν-Mediated Phagocytosis of Apoptotic Cells in Drosophila Embryos *’. Journal of Biological Chemistry 286 (29): 25770–77. 10.1074/jbc.M110.204503.

Pais, Inês S., Rita S. Valente, Marta Sporniak, and Luis Teixeira. 2018a. ‘Drosophila Melanogaster Establishes a Species-Specific Mutualistic Interaction with Stable Gut-Colonizing Bacteria’. PLOS Biology 16 (7): e2005710. 10.1371/journal.pbio.2005710.

Pais, Inês S., Rita S. Valente, Marta Sporniak, and Luis Teixeira. 2018b. ‘Drosophila Melanogaster Establishes a Species-Specific Mutualistic Interaction with Stable Gut-Colonizing Bacteria’. PLOS Biology 16 (7): e2005710. 10.1371/journal.pbio.2005710.

Pearson, Alan M., Katalin Baksa, Mika Rämet, Meredith Protas, Mary McKee, Dennis Brown, and R. Alan B. Ezekowitz. 2003. ‘Identification of Cytoskeletal Regulatory Proteins Required for Efficient Phagocytosis in Drosophila’. Microbes and Infection 5 (10): 815–24. 10.1016/s1286-4579(03)00157-6.

Petrignani, Bianca, Samuel Rommelaere, Ketty Hakim-Mishnaevski, Florent Masson, Elodie Ramond, Reut Hilu-Dadia, Mickael Poidevin, Shu Kondo, Estee Kurant, and Bruno Lemaitre. 2021. ‘A Secreted Factor NimrodB4 Promotes the Elimination of Apoptotic Corpses by Phagocytes in Drosophila’. EMBO Reports 22 (9): e52262. 10.15252/embr.202052262.

Ramond, Elodie, Bianca Petrignani, Jan Paul Dudzic, Jean-Philippe Boquete, Mickaël Poidevin, Shu Kondo, and Bruno Lemaitre. 2020. ‘The Adipokine NimrodB5 Regulates Peripheral Hematopoiesis in Drosophila’. The FEBS Journal 287 (16): 3399–3426. 10.1111/febs.15237.

Rommelaere, Samuel, Jean-Philippe Boquete, Jérémie Piton, Shu Kondo, and Bruno Lemaitre. 2019. ‘The Exchangeable Apolipoprotein Nplp2 Sustains Lipid Flow and Heat Acclimation in *Drosophila*’. Cell Reports 27 (3): 886–899.e6. 10.1016/j.celrep.2019.03.074.

Sarov, Mihail, Christiane Barz, Helena Jambor, Marco Y Hein, Christopher Schmied, Dana Suchold, Bettina Stender, et al. 2016. ‘A Genome-Wide Resource for the Analysis of Protein Localisation in Drosophila’. Edited by Hugo J Bellen. eLife 5 (February):e12068. 10.7554/eLife.12068.

Segawa, Katsumori, Sachiko Kurata, Yuichi Yanagihashi, Thijn R. Brummelkamp, Fumihiko Matsuda, and Shigekazu Nagata. 2014. ‘Caspase-Mediated Cleavage of Phospholipid Flippase for Apoptotic Phosphatidylserine Exposure’. *Science (New York*, N.Y.) 344 (6188): 1164–68. 10.1126/science.1252809.

Segawa, Katsumori, and Shigekazu Nagata. 2015. ‘An Apoptotic “Eat Me” Signal: Phosphatidylserine Exposure’. Trends in Cell Biology 25 (11): 639–50. 10.1016/j.tcb.2015.08.003.

Serizier, Sandy B., and Kimberly McCall. 2017. ‘Scrambled Eggs: Apoptotic Cell Clearance by Non-Professional Phagocytes in the Drosophila Ovary’. Frontiers in Immunology 8:1642. 10.3389/fimmu.2017.01642.

Shklover, Jeny, Flonia Levy-Adam, and Estee Kurant. 2015. ‘Apoptotic Cell Clearance in Development’. Current Topics in Developmental Biology 114:297–334. 10.1016/bs.ctdb.2015.07.024.

Shklyar, Boris, Flonia Levy-Adam, Ketty Mishnaevski, and Estee Kurant. 2013. ‘Caspase Activity Is Required for Engulfment of Apoptotic Cells’. Molecular and Cellular Biology 33 (16): 3191–3201. 10.1128/MCB.00233-13.

Sinenko, Sergey A., and Bernard Mathey-Prevot. 2004. ‘Increased Expression of Drosophila Tetraspanin, Tsp68C, Suppresses the Abnormal Proliferation of Ytr-Deficient and Ras/Raf-Activated Hemocytes’. Oncogene 23 (56): 9120–28. 10.1038/sj.onc.1208156.

Somogyi, Kálmán, Botond Sipos, Zsolt Pénzes, Eva Kurucz, János Zsámboki, Dan Hultmark, and István Andó. 2008. ‘Evolution of Genes and Repeats in the Nimrod Superfamily’. Molecular Biology and Evolution 25 (11): 2337–47. 10.1093/molbev/msn180.

Stephenson, Holly N., Robert Streeck, Florian Grüblinger, Christian Goosmann, and Alf Herzig. 2022. ‘Hemocytes Are Essential for Drosophila Melanogaster Post-Embryonic Development, Independent of Control of the Microbiota’. Development (Cambridge, England) 149 (18): dev200286. 10.1242/dev.200286.

Tollis, Sylvain, Anna E. Dart, George Tzircotis, and Robert G. Endres. 2010. ‘The Zipper Mechanism in Phagocytosis: Energetic Requirements and Variability in Phagocytic Cup Shape’. BMC Systems Biology 4 (1): 149. 10.1186/1752-0509-4-149.

Troha, Katia, Peter Nagy, Andrew Pivovar, Brian P. Lazzaro, Paul S. Hartley, and Nicolas Buchon. 2019. ‘Nephrocytes Remove Microbiota-Derived Peptidoglycan from Systemic Circulation to Maintain Immune Homeostasis’. Immunity 51 (4): 625–637.e3. 10.1016/j.immuni.2019.08.020.

Tung, Tran Thanh, Kaz Nagaosa, Yu Fujita, Asana Kita, Hiroki Mori, Ryo Okada, Saori Nonaka, and Yoshinobu Nakanishi. 2013. ‘Phosphatidylserine Recognition and Induction of Apoptotic Cell Clearance by Drosophila Engulfment Receptor Draper’. The Journal of Biochemistry 153 (5): 483–91. 10.1093/jb/mvt014.

Ulvila, Johanna, Leena-Maija Vanha-Aho, and Mika Rämet. 2011. ‘Drosophila Phagocytosis – Still Many Unknowns under the Surface’. APMIS 119 (10): 651–62. 10.1111/j.1600-0463.2011.02792.x.

Uribe-Querol, Eileen, and Carlos Rosales. 2020. ‘Phagocytosis: Our Current Understanding of a Universal Biological Process’. Frontiers in Immunology 11:1066. 10.3389/fimmu.2020.01066.

Yamaguchi, Hiroshi, Takashi Fujimoto, Shinobu Nakamura, Koichiro Ohmura, Tsuneyo Mimori, Fumihiko Matsuda, and Shigekazu Nagata. 2010. ‘Aberrant Splicing of the Milk Fat Globule-EGF Factor 8 (MFG-E8) Gene in Human Systemic Lupus Erythematosus’. European Journal of Immunology 40 (6): 1778–85. 10.1002/eji.200940096.

Zhang, Yang, and Jeffrey Skolnick. 2005. ‘TM-Align: A Protein Structure Alignment Algorithm Based on the TM-Score’. Nucleic Acids Research 33 (7): 2302–9. 10.1093/nar/gki524.

