## Supplementary document for "The secreted Nimrod NimB1 negatively regulates early steps of apoptotic cell phagocytosis in *Drosophila*"

### Supplementary documents:

**Supplementary document S1: NimB1 localization and binding to ACs**

(A-D) Representative confocal microscopy images of macrophages dissected from *Hml^p2a^> UAS-SP^vkg^-RFP* third instar larvae (A, negative control) and *Hml^p2a^ >UAS-NimB1-RFP* (B-D) larvae, incubated with ACs labeled with a far-red Cell Trace^TM^ dye (AC FRED, purple). RFP signal marks NimB1 localization. Differential interference contrast (DIC) imaging is shown for cell morphology. Scale bars = 10 μm for all panels.

(A) Macrophages expressing the control secreted protein (*UAS-SP^vkg^-RFP*) exhibit diffuse RFP signal with no specific binding to ACs.

(B) Macrophages expressing NimB1-RFP show distinct patch-like RFP localization of the protein in the cytosolic region of macrophages (arrows) and localized on the edges of the cell membrane (arrowheads). 3D reconstructions done with the Imaris Viewer 10.2.0. software

(C) Tilted image of (B)

(D) Zoomed view of (C) showing clustering of NimB1-RFP at the cell membranes of macrophages.

(E) Quantification of proportion of macrophages with membrane-associated RFP signal from *Hml^p2a^>UAS-SP^vkg^-RFP* (control) and *Hml^p2a^>UAS-NimB1-RFP*. Data are presented as the mean ± SD from three independent experiments with at least 50 cells analyzed per experiment. (*****P* < 0.0001, by unpaired *t-test*).

(F) Fluorescence analysis of nephrocytes in control larvae *Lpp^ts^* >*w^1118^* (left panel) and larvae expressing *Lpp^ts^* >*UAS-NimB1-RFP* (right panel). Images were acquired via whole-larvae microscopy. Scale bar = 500 μm.

**Supplementary document S2: Description and validation of *NimB1* and *NimB1*, *NimB4* double mutants**

(A) Schematic representation of the *NimB* locus, highlighting targeted genetic modifications. The *NimB* gene cluster (2L:13,943,501–13,972,167) includes *NimB1–5*. CRISPR-Cas9 editing generated the *NimB1*^229^ deletion allele by removing an 11 bp sequence in exon 1, resulting in a frameshift mutation. The *NimB1^crimic^* mutant was generated by the insertion of a *T{CRIMIC.TG4.2}* transgene into intron 1 of *NimB1*

*.* A double mutant (*NimB1*^61^, *NimB4*^sk2^) was created by targeting *NimB1* in a *NimB4*^sk2^ background. CRISPR-Cas9 editing resulted in generated a 13 bp deletion in exon 1 of *NimB1* in the background on *NimB4^sk2^* featuring a 14 bp deletion.

(B) Quantification of *NimB1* overexpression levels with hemocyte-specific *Hml^p2a^-Gal4* driver in various genotypes using RT-qPCR, normalized to *Rp49*. Data are presented as mean ± SD from three independent experiments. (*P < 0.05, **P < 0.01, by one-way ANOVA with Tukey’s post hoc test).

(C) Western blot analysis of hemolymph extracts from third instar larvae expressing *UAS-SP^vkg^-RFP* (control) or *UAS-NimB1-RFP* under the *Lpp^ts^* driver. A 70 kDa band confirms NimB1-RFP secretion into the hemolymph, while a 25 kDa band corresponds to RFP protein.

**Supplementary document S3: Impact of NimB1 on hemocyte count and shape**

(A) Hemocyte counts from hemolymph samples of five third instar larvae per genotype, quantified using FACS analysis. Data are derived from three independent experiments and represented as individual data points with mean ± SD. (*****P* < 0.0001, by one-way ANOVA with Tukey’s post hoc test).

(B) Quantification of hemocyte cell area based on confocal images in (D). Data are derived from three independent experiments and represented as individual data points with means ± SD. (*****P* < 0.0001, by one-way ANOVA with Tukey’s post hoc test).

(C) Filopodia-to-lamellipodia ratio per frame across genotypes, indicating cytoskeletal dynamics and remodeling in macrophages. Data are derived from three independent experiments and represented as individual data points with means ± SD. (*****P* < 0.0001, by one-way ANOVA with Tukey’s post hoc test).

(D) Representative confocal images of hemocyte adhesion and spreading, visualized with Phalloidin (cyan) across genotypes: *w^1118^*, *NimB1^crimic^*, *NimB4^sk2^ and NimB1^61^,NimB4^sk2^*.

**Supplementary document S4: *Ex vivo* phagocytic assay of ACs at 1-hour and 2-hour time points. Idem Figure 4.**

(A) Phagocytic activity measured after 1-hour incubation of CFSE-labeled ACs with macrophages isolated from wild-type *(w^1118^), NimB1^crimic^, NimB1^229/crimic^, NimB1^61^, NimB4^sk2^, NimB4^sk2^,* and *draper^Δ5^* larvae. Phagocytic activity is assessed by the phagocytic index (left panel), calculated as the product of the mean fluorescence intensity (MFI, middle panel) multiplied by the percentage of macrophages actively phagocytosing ACs (right panel). Data are derived from three independent experiments and represented as individual data points with mean ± SD.

(B) Phagocytic activity measured after 2-hour incubation under the same conditions as in (A).

Statistical analysis: Data are derived from three independent experiments and represented as individual data points with means ± SD. (*p < 0.05, **p < 0.01, ***p < 0.001, **p < 0.0001, ns = not significant, by one-way ANOVA with post hoc Tukey’s multiple comparisons test).

**Supplementary document S5: Rescue of phagocytic activity in *NimB1^crimic^* mutants. Idem Figure 4.**

(A) Phagocytic activity measured after 1-hour incubation of CFSE-labeled ACs with macrophages isolated from wild-type (*w^1118^*), *NimB1^crimic^*, *NimB1^crimic^* + *UAS-NimB1*, *NimB1^crimic^* + *UAS*-*NimB1-RFP*, and *draper^Δ5^* larvae. Phagocytic activity is assessed by the phagocytic index (left panel), calculated as the product of the mean fluorescence intensity (MFI, middle panel) and the percentage of macrophages actively phagocytosing apoptotic cells (right panel).

(B) Phagocytic activity measured after 2-hour incubation under the same conditions as in (A).

Statistical analysis: Data are derived from three independent experiments and represented as individual data points with means ± SD. (*p < 0.05, **p < 0.01, ***p < 0.001, **p < 0.0001, ns = not significant, by one-way ANOVA with post hoc Tukey’s multiple comparisons test).

**Supplementary document S6: Phagocytic assay for bacterial uptake by *NimB1*** macrophages

(A) Phagocytic index of macrophages isolated from wild-type (*w^1118^*), *NimB1^crimic^*, *NimB1^61^*,*NimB^sk2^*, and *NimC1;eater^1^* double mutant larvae incubated with *E. coli* for 1 hour.

(B) Phagocytic index of macrophages from the same genotypes as in (A) incubated with *S. aureus* for 1 hour.

Statistical analysis: Data are derived from three independent experiments and represented as individual data points with means ± SD. (**p < 0.01, ns = not significant, by one-way ANOVA with post hoc Tukey’s multiple comparisons test).
