## Supplementary figures for "The secreted Nimrod NimB1 negatively regulates early steps of apoptotic cell phagocytosis in *Drosophila*"

### Supplementary documents

### Supplementary document S1

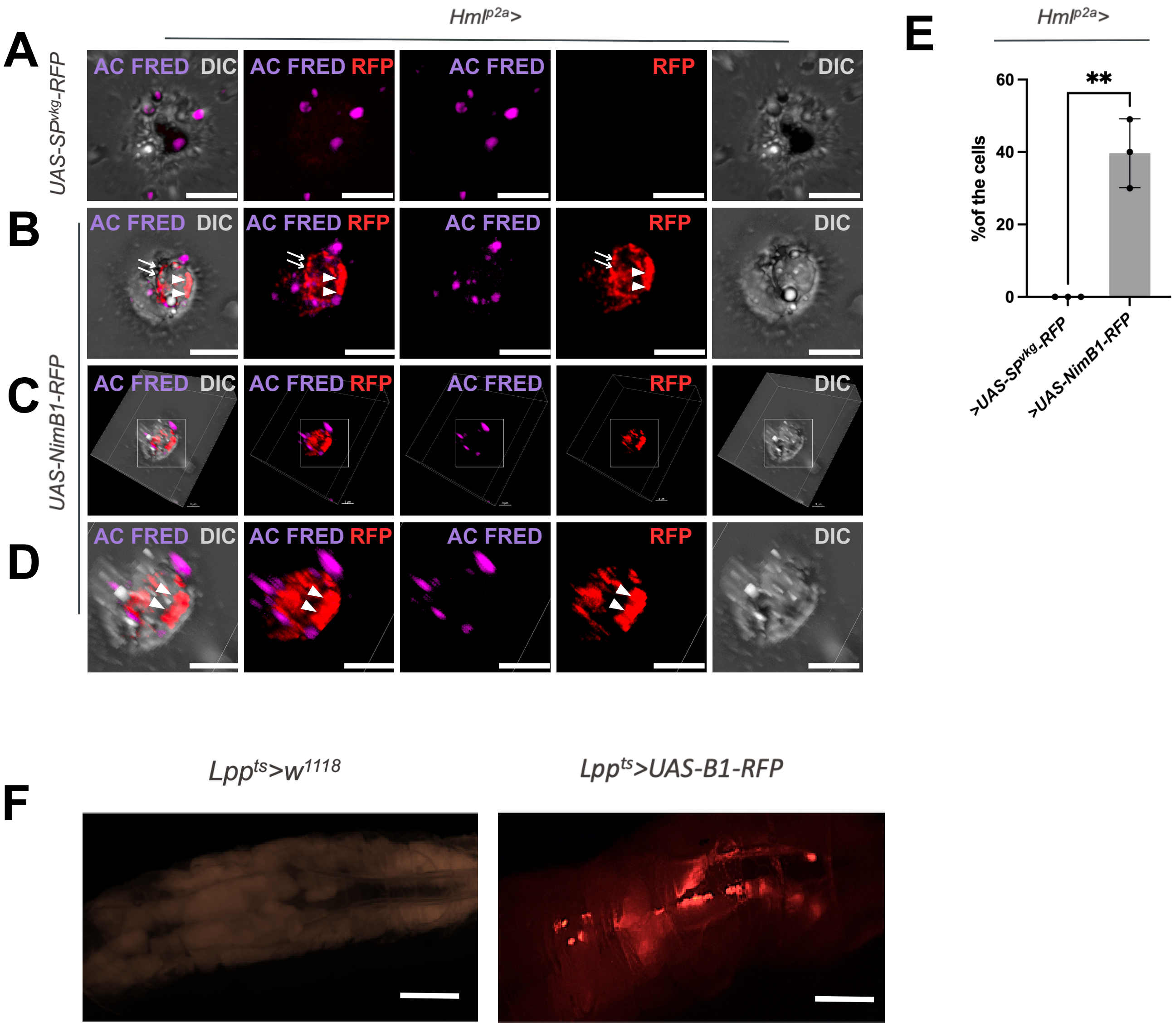

### Supplementary document S2

**A**

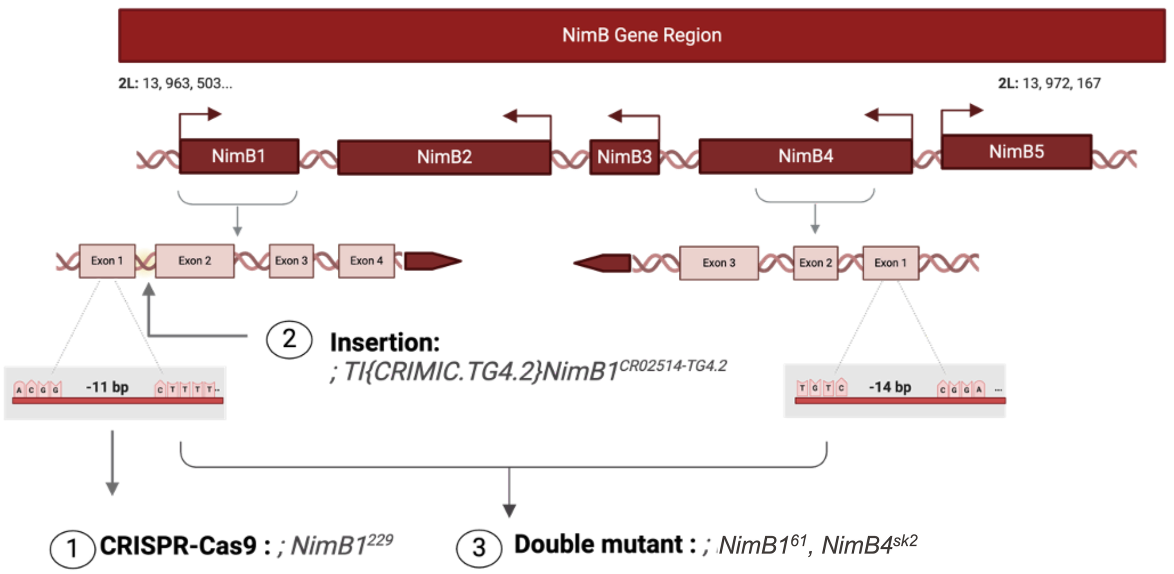


**B**

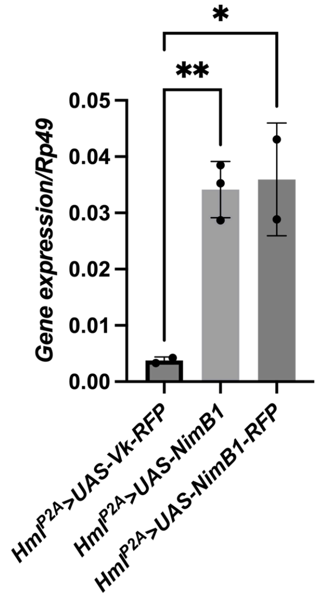


**C**

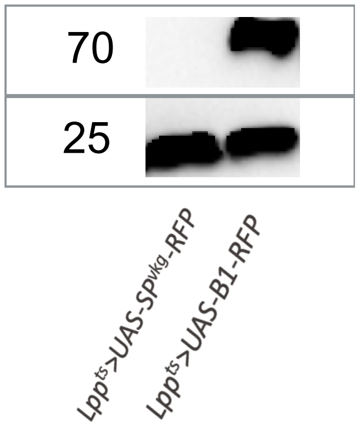

### Supplementary document S3

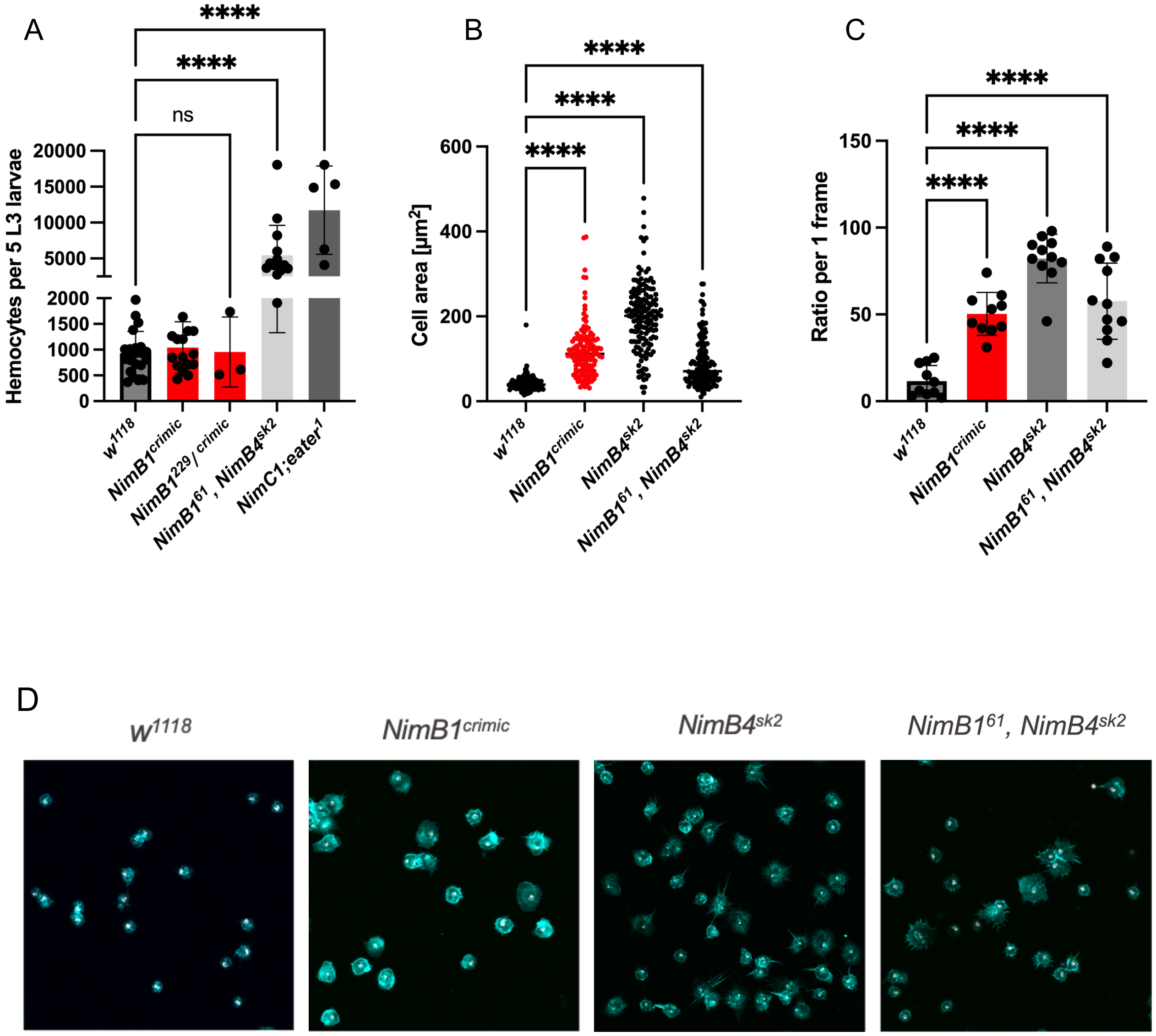

### Supplementary document S4

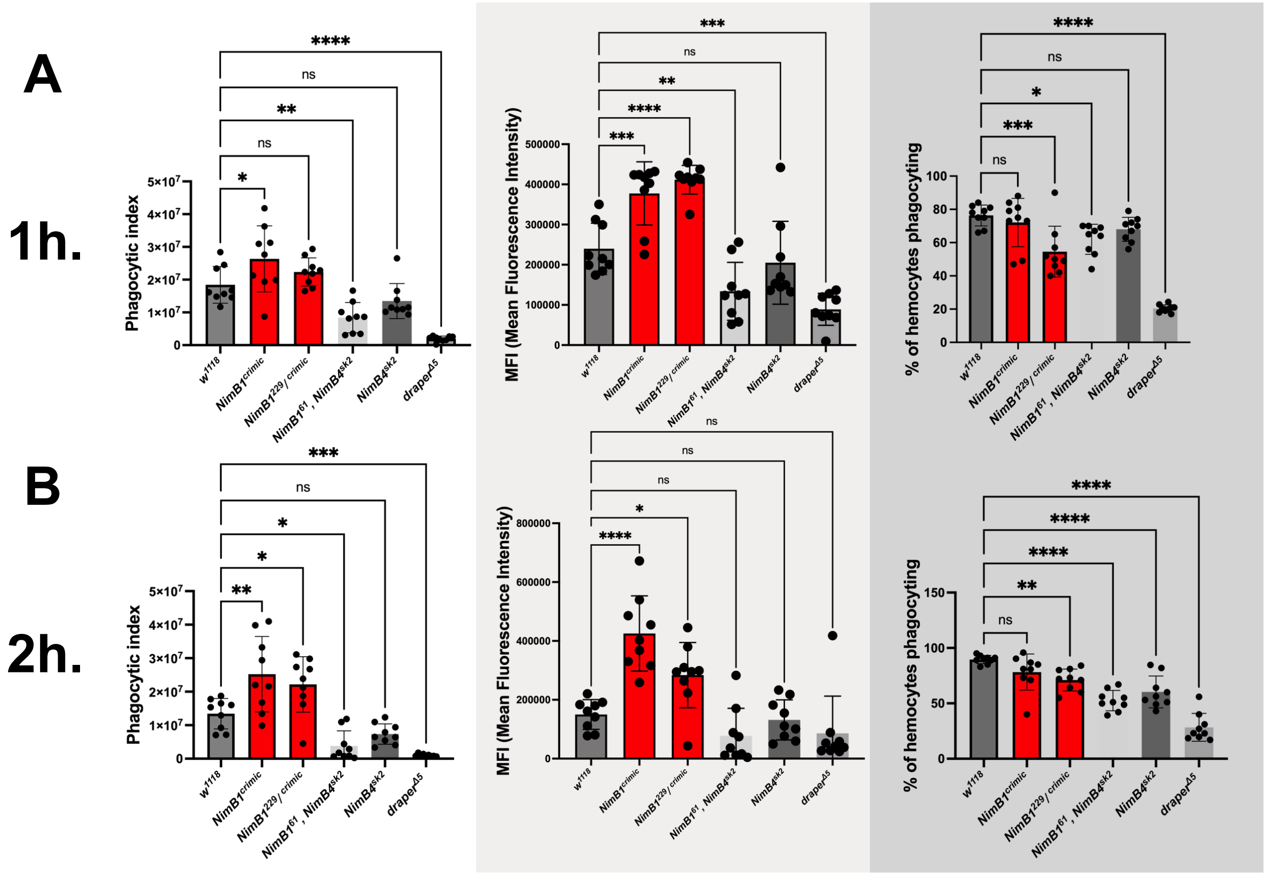

### Supplementary document S5

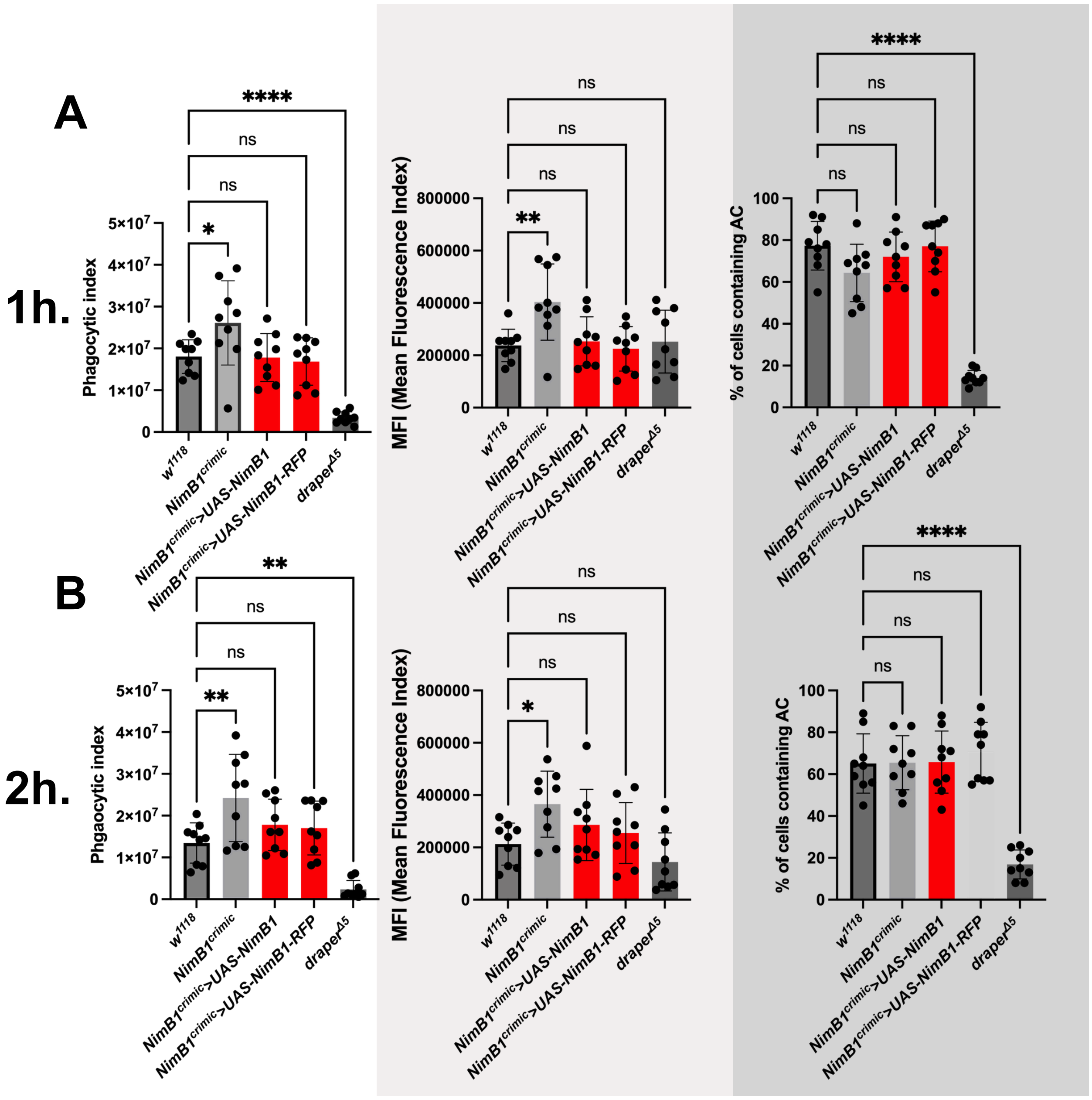

### Supplementary document S6

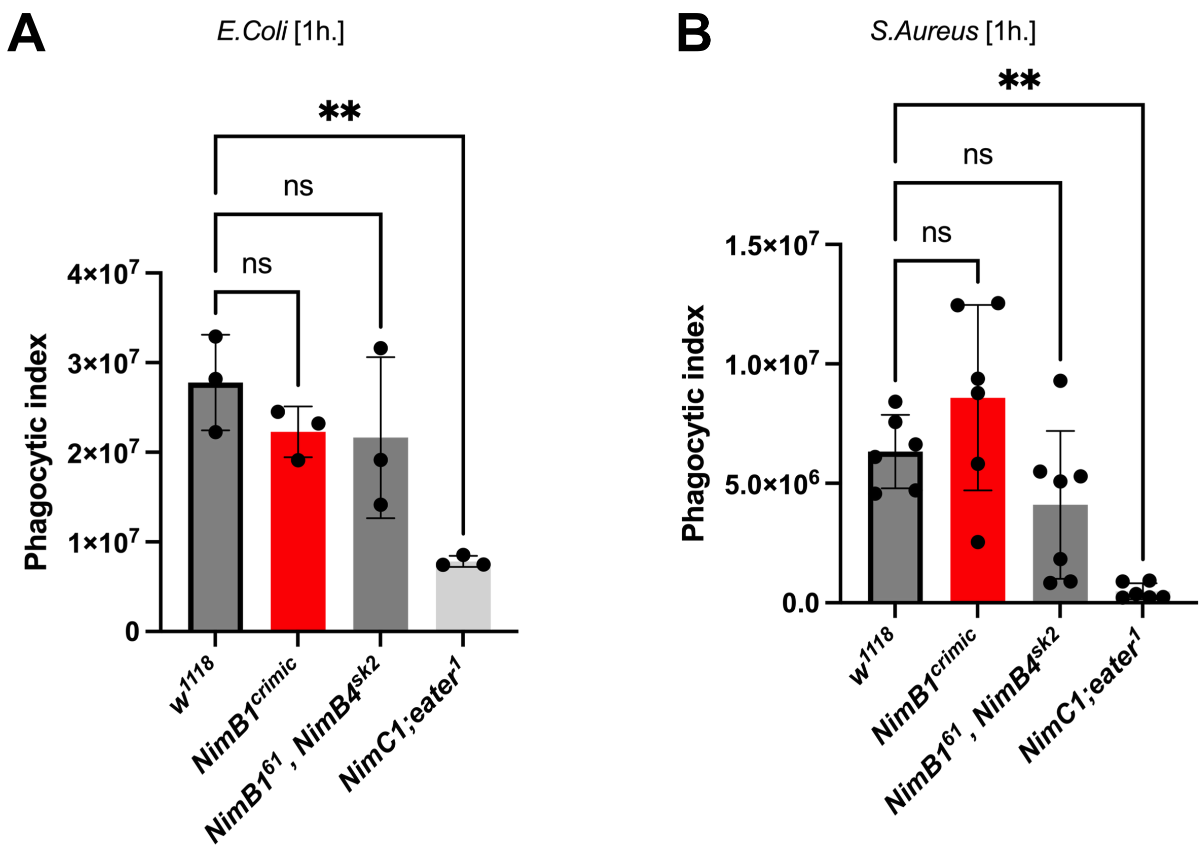
